# Time-resolved transcriptomics reveal a mechanism of host niche defense: beneficial root endophytes deploy a host-protective antimicrobial GH18-CBM5 chitinase

**DOI:** 10.1101/2023.12.29.572992

**Authors:** Ruben Eichfeld, Lisa K. Mahdi, Concetta De Quattro, Laura Armbruster, Asmamaw B. Endeshaw, Shingo Miyauchi, Margareta J. Hellmann, Stefan Cord-Landwehr, Igor Grigoriev, Daniel Peterson, Vasanth Singan, Kathleen Lail, Emily Savage, Vivian Ng, Gregor Langen, Bruno M. Moerschbacher, Alga Zuccaro

## Abstract

Associations between plants and beneficial root-endophytic fungi enhance plant performance by improving nutrient uptake, abiotic stress tolerance and disease resistance. To successfully colonize different host plants and defend their host niche against competing microbes, but also to cooperate with beneficial bacterial members of the microbiota, root endophytes such as Sebacinales secrete a multitude of tightly regulated effector-proteins and carbohydrate-active enzymes. However, the functions, specificity, and regulation of these proteins remain poorly understood. In this study, we employ time-resolved transcriptomics to analyse the gene expression profiles of two Sebacinales members interacting with organisms from different kingdoms of life. We identified crucial genes for plant colonization and intermicrobial competition, including a fungal GH18-CBM5 chitinase specifically upregulated in response to the phytopathogenic fungus *Bipolaris sorokiniana*. This chitinase protects the plant hosts against the pathogen, reducing fungal biomass and disease symptoms in barley and *Arabidopsis thaliana*. Our findings shed light on interaction partner specific gene expression in Sebacinales endophytes, with potential applications in enhancing plant health and resilience.

**Bullet points:**

- Both *Serendipita indica* (*Si*) and *Serendipita vermifera* (*Sv*) show similar transcriptional responses to three host species and the phytopathogen *Bipolaris sorokiniana* (*Bs*), indicating common interaction principles between Sebacinales and plant hosts or fungi.
- These shared mechanisms involve the activation of effector genes like small secreted proteins and carbohydrate-active enzymes.
- Cooperation with beneficial bacteria elicits only minimal transcriptomic alterations in Sebacinales compared to plants and *Bs*.
- Sebacinales respond to *Bs* by upregulating a specific GH18-CBM5 chitinase unique to Basidiomycota within the fungal kingdom, inhibiting *Bs* growth and reducing disease symptoms in *Arabidopsis thaliana* and barley.

## Introduction

Beneficial root-endophytic fungi are major players within the consortia of plant-associated microorganisms collectively referred to as “plant microbiome” (Glynou *et al*., 2016; Glynou *et al*., 2018; Trivedi *et al*., 2020; Mahdi *et al*., 2022). While the composition of this microbiome varies between different host plants and depends on environmental factors (Tkacz *et al*., 2015; Strullu-Derrien *et al*., 2018), a balanced microbiome contributes to plant health and productivity by improving host nutrient uptake and increasing the resistance to biotic and abiotic stress (Raaijmakers *et al*., 2009; Hermosa *et al*., 2012; Finkel *et al*., 2017; Mahdi *et al*., 2022). Such beneficial properties have been found in plant interactions with ectomycorrhizal (ECM) and arbuscular mycorrhizal (AM) fungi, as well as certain fungal endophytes (Zuccaro *et al*., 2014). Symbiotic interactions have evolved over millions of years, giving rise to fine-tuned relationships not only between microbes and their host plants but also among the diverse constituents of the microbiome themselves (Mesny *et al*., 2023). The health of plants is directly influenced by these intermicrobial relationships. This is illustrated by microorganisms which manifest high pathogenic potential in mono-associations but are effectively restrained in a microbial community context (Sarkar *et al*., 2019; Mesny *et al*., 2021; Mahdi *et al*., 2022). The underlying mechanism behind this phenomenon is intermicrobial competition and cooperation, as members of the microbiome have evolved strategies to defend their host niche against phytopathogenic intruders. The beneficial root endophytic fungus *Serendipita vermifera* for instance acts synergistically with bacterial microbiome members to protect barley against the phytopathogenic fungus *Bipolaris sorokiniana* (Mahdi *et al*., 2022). *B. sorokiniana* causes diseases in cereals, including common root rot and spot blotch, resulting in dramatic yield losses especially in warmer agricultural areas (Kumar *et al*., 2002). While some fungi outcompete other microbes by resource sequestration or inducing host immunity, *S. vermifera* is thought to antagonize *B. sorokiniana* mostly directly through the secretion of effectors and hydrolytic enzymes, thus reducing the virulence potential of the plant pathogen prior to host colonization (Sarkar *et al*., 2019). Effectors were originally described as secreted proteins which suppress plant immunity in order to promote microbial colonization and reproduction (De Wit *et al*., 2009). It has been speculated that the broad host range of Sebacinales is a consequence of their expanded repertoire of effector proteins. Indeed, the genomes of *S. vermifera* and the closely related *S. indica* encompass an arsenal of genes encoding for proteins involved in carbohydrate binding, plant cell wall degradation and protein hydrolysis, as well as numerous small-secreted proteins (SSPs) with effector-like properties (Zuccaro *et al*., 2011; Zuccaro *et al*., 2014).

Recent discoveries in phytopathogenic fungi have provided first insights into the importance of fungal effectors during intermicrobial competition (Veneault-Fourrey & Martin, 2011; Hemetsberger *et al*., 2012; Win *et al*., 2012; Lo Presti *et al*., 2015; Snelders *et al*., 2022). The soil-borne fungus *Verticillium dahliae*, for instance, secretes the virulence effector and antimicrobial protein *Vd*Ave1, which suppresses antagonistic bacteria and facilitates the infection of tomato plants (Snelders *et al*., 2023). Transcriptomic analyses of *S. vermifera* and *B. sorokiniana* during bipartite confrontation in soil and tripartite interactions with barley revealed that *S. vermifera* and *B. sorokiniana* express unique sets of effectors during intermicrobial confrontation or root colonization. In addition, a group of core effectors was induced during both fungal confrontation and host colonization (Sarkar *et al*., 2019). The extent to which this finding can be extrapolated to other fungi and host plants remains unclear. Furthermore, it is uncertain whether the endophyte *S. vermifera*, as previously observed in the case of the plant pathogen *V. dahliae* (Snelders *et al*., 2023), expresses microbe-targeting effectors that modulate members of the microbiome, including root-associated bacteria and fungi.

In this study, we conducted extensive time-resolved transcriptomic analysis in *S. indica* and *S. vermifera*, examining their transcriptional responses when exposed to monocot and dicot host plants, the phytopathogen *B. sorokiniana*, or beneficial root-associated bacteria. Our focus was on investigating putative effector genes, particularly small-secreted proteins (SSPs) and carbohydrate-active enzymes (CAZymes). We revealed that while most Sebacinales effectors are triggered by both host plants and plant-associated microbes, specific subsets are exclusive to either the host or microbe. Notably, the expression of microbe-specific effectors is predominantly triggered in the presence of *B. sorokiniana* but not the beneficial root-associated bacteria. Additionally, we characterized a GH18-CBM5 chitinase in *S. indica* and *S. vermifera*, exclusively induced in response to *B. sorokiniana*. These chitinases are unique to Basidiomycota within the fungal kingdom. They possess catalytic activity, effectively inhibiting spore germination and growth of *B. sorokiniana*, thereby shielding barley and *Arabidopsis thaliana* from the phytopathogenic fungus. Thus, the GH18-CBM5 serves as a pivotal antimicrobial effector for safeguarding the root niche. Understanding how beneficial fungi utilize effectors to establish themselves within host plants and safeguard their root niche against competing microorganisms and phytopathogens will deepen our knowledge of complex multiorganismal interactions and lead to the development of biocontrol strategies aimed at reducing reliance on chemical pesticides in agriculture.

## Results

### Co-culture with root-associated bacteria elicits only minimal changes in gene transcription patterns in beneficial endophytes

To investigate the molecular mechanisms of how *S. indica* (*Si*) and *S. vermifera* (*Sv*) interact with a wide range of organisms from different kingdoms, we generated an RNA-seq dataset covering bipartite interactions of *Si* or *Sv* with the plant hosts *Hordeum vulgare* (*Hv*), *Brachypodium distachyon* (*Bd*) and *A. thaliana* (*At*), as well as with the plant pathogen *B. sorokiniana* (*Bs*) or a bacterial synthetic community (SynCom) consisting of four taxonomically distinct bacteria derived from *At* roots (Mahdi *et al*., 2022). We addressed possible temporal differences in the establishment of interaction stages by including samples collected at four different time points post inoculation (Figure 1A). To assess the similarity between the different treatments, we conducted a principal component analysis. We found that the transcriptional profiles of *Si* and *Sv* separated into three distinct groups based on their interaction partners: plant hosts (green), the bacterial SynCom (red), and the phytopathogenic fungus *Bs* (orange) (Figure 1B).

**Figure 1:**
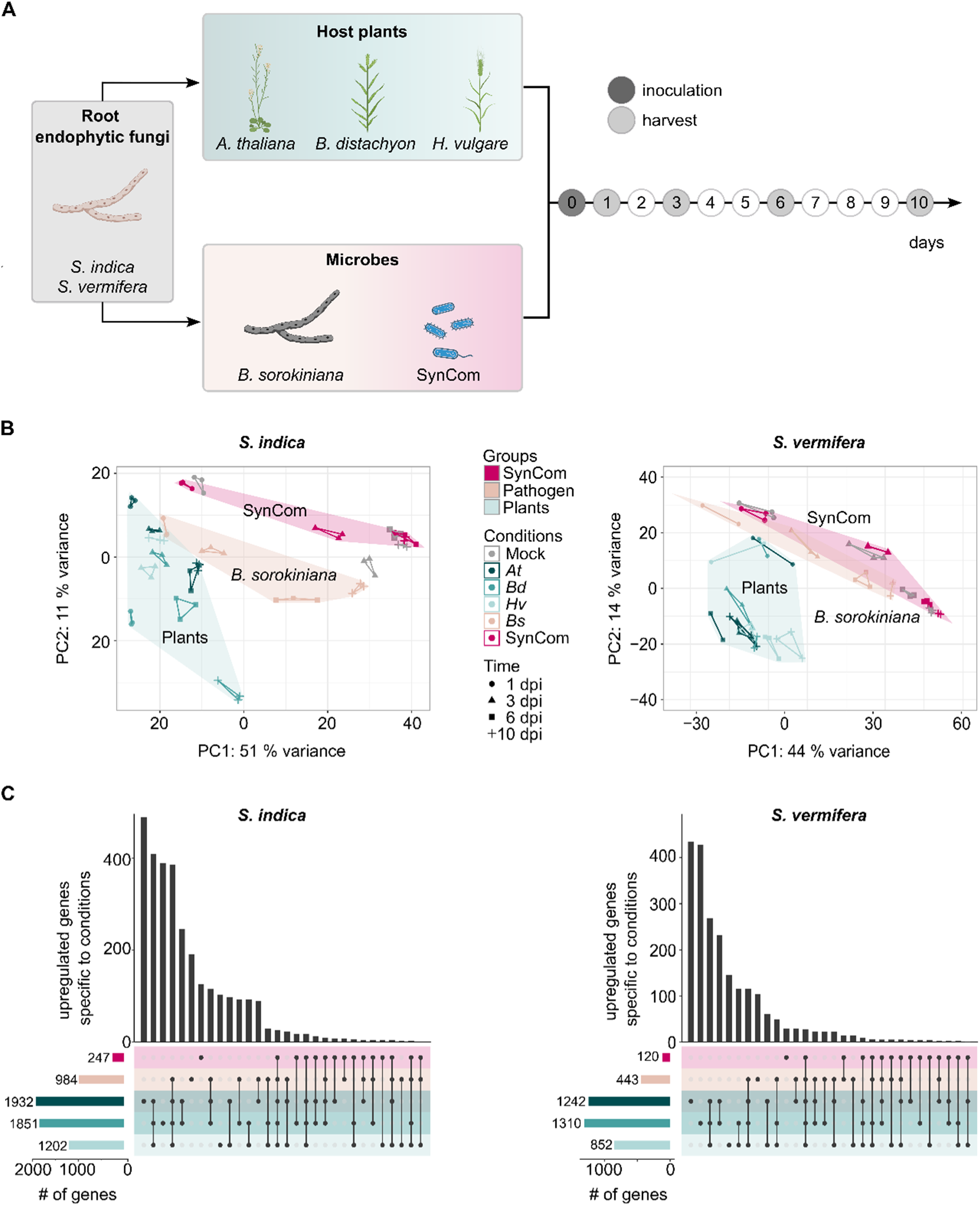
Transcriptional response of *S. indica* and *S. vermifera* to different interaction partners. **A)** Schematic overview of the experimental setup. Bipartite interactions between *S. indica* (*Si*) or *S. vermifera* (*Sv*) and the plant hosts *Arabidopsis thaliana (At), Brachypodium distachyon (Bd)* and *Hordeum vulgare (Hv)* or the microbes *Bipolaris sorokiniana* (*Bs*) and a synthetic bacterial community (SynCom) at different days past inoculation (dpi). **B)** PCA plots comprising the top 500 most variable genes of *Si* (left) and *Sv* (right) in response to the different interaction partners across all time points. Each biological replicate is shown individually and connected with lines to the respective other replicates of the same treatment and time point. Transcriptomic responses to host plants, the SynCom and the plant pathogen *Bs* are highlighted with green, dark red and orange backgrounds, respectively. **C)** UpSet plot of upregulated genes (FDR-adjusted p-value < 0.05 and log_2_FC > 2) aggregated across all time points in *Si* (left) and *Sv* (right) in response to *At*, *Bd*, *Hv*, *Bs* and the SynCom.

To further investigate the gene expression changes induced in *Si* and *Sv* during biotic interactions, we performed a differential gene expression analysis. When comparing transcriptional patters between axenically cultured fungi and fungi challenged with hosts or microbes, we found a total of 4,838 (*Si*) or 5,606 (*Sv*) genes which were differentially expressed (>2 log_2_FC or <-2 log_2_FC, adjusted p-value <0.05) in at least one of the interactions at one or more timepoints. These differentially expressed genes (DEGs) accounted for approximately 40 % of annotated *Si* and 37 % of annotated *Sv* genes. We focussed our analysis on the upregulated genes with a log_2_FC >2 (2,999 *Si* and 2,185 *Sv* genes). To identify commonalities and differences in gene expression during the interaction with plant hosts and microbes, we collapsed the significantly upregulated genes at different time points for each treatment (Figure 1C). While both *Si* and *Sv* responded to all plant hosts and to the plant pathogen *Bs* with extensive transcriptional alterations, bacteria did not elicit pronounced transcriptional changes in the beneficial root endophytes.

### Sebacinales express a core set of genes in response to monocot and dicot hosts

The responses of *Si* and *Sv* to the three plant hosts largely overlapped, with 837 (*Si*) and 393 (*Sv*) genes upregulated in the presence of all three hosts. These accounted for 31 % and 19 % of all 2,676 (*Si*) or 2,038 (*Sv*) plant-inducible genes, respectively (Supplementary Table S1). This suggests that exposure of *Si* and *Sv* to different plant species triggered the expression of a set of core genes required for host colonization in both monocots and dicots. These genes included the nucleotidase *E5’NT* (Pirin1_71782; PIIN_01005 for *Si* and Sebve1_ 17804 for *Sv*) and the nuclease *NucA* (Pirin1_72917; PIIN_02121 for *Si* and Sebve1_52856 for *Sv*), which are involved in suppression of immunity and initiation of host cell death *via* the production of small active molecules (Nizam *et al*., 2019; Dunken *et al*., 2022). Restricted host cell death is important for successful colonization of plant hosts by the two endophytes and is considered a nutritional strategy of Sebacinales, which have retained the saprotrophic capabilities of their ancestors (Deshmukh *et al*., 2006; Qiang *et al*., 2012). Intracellular colonization of all three plant hosts was associated with upregulation of fungal proteases and CAZymes, which can degrade host-derived proteins as well as plant cell walls and may serve to convey entry into the host cell as well as a source of nitrogen and carbon for the endophytes. An organic nitrogen source is particularly relevant for *Si*, as this fungus seems to be unable to utilize nitrate as a nitrogen source (Olivieri *et al*., 2002; Naumann *et al*., 2011; Zuccaro *et al*., 2011; Lahrmann & Zuccaro, 2012; Balestrini *et al*., 2014; Zuccaro *et al*., 2014; Jashni *et al*., 2015a; Jashni *et al*., 2015b; Tang *et al*., 2021; Valadares *et al*., 2021). Interestingly, we found genes encoding glycosyltransferase enzymes with GT32 (Pirin1_74046; PIIN_03242 in *Si* and Sebve1_24669 in *Sv*) and GT8 domains (Pirin1_77790; PIIN_06984 in *Si* and Sebve1_26577 in *Sv*) that were upregulated at later time points in all plant hosts. These enzymes may be involved in fungal carbohydrate metabolism required for the establishment of successful long-lasting symbioses and were not responsive to microbes.

Despite commonalities between responses across all plant hosts, significant sets of genes were exclusively induced in the presence of a specific plant species. Several of these host-species specific genes seemed to serve similar functions; for instance, Pirin1_74456 (PIIN_03655; upregulated specifically in response to *At*) and Pirin1_80981 (PIIN_10163, upregulated specifically in response to *Bd*), both of which encode CE4 polysaccharide deacetylases. These deacetylases can be exploited by root endophytes to modulate chitin in their cell walls, aiding in evading plant immunity. Moreover, we have pinpointed enzymes that hydrolyse specific substrates present in monocots but absent in *At* (Supplementary Figure 1 and 2), including the previously identified GH10 and GH11 family xylanases and AA9 family polysaccharide monooxygenases (Lahrmann *et al*., 2013).

### Endophytes display conserved transcriptional responses to plant hosts and the phytopathogenic fungus *Bs*

To gain more insight into the biological functions of the Sebacinales genes induced in response to plant hosts or microbes, we analysed the two sets of genes separately (Supplementary Figure 3). Employing K-means analysis, we divided both sets into three distinct clusters, representing genes upregulated either throughout the interaction, in early or in late stages of interaction. A gene ontology (GO) term analysis revealed that similar processes were induced in response to microbes and plants in both Sebacinales. These include “carbohydrate metabolic process” (GO:0005975), “proteolysis” (GO:0006508) and “transport” (GO:0006810). Genes assigned to all three terms were induced during all stages of colonization and might relate to nutritional processes, indicating that *Si* and *Sv* take up nutrients from plant as well as microbial biomatter. Another GO term likely related to nutrient acquisition was “cell wall catabolic process” (GO:0016998). Interestingly, genes related to this term were strongly induced in both Sebacinales in the early phases of the response to *Bs*, but not plants. The induction of this specific set of genes could be interpreted as a sign of mycoparasitism.

Besides a large overlap of upregulated genes between all plant hosts and specific genes induced in response to *Bs*, we identified a set of commonly upregulated genes in response to plants and *Bs* (787 genes in *Si*; 291 genes in *Sv*). These genes accounted for 80 and 66 % of the total *Bs* inducible genes in *Si* and *Sv*, suggesting underlying mechanistic parallels in the interaction of Sebacinales with plants and other fungi. An example for a gene which was induced upon exposure of Sebacinales to plant hosts and the plant pathogen *Bs* is *WSC3* (Pirin1_76632; PIIN_05825 in *Si* and Sebve1_309621 in *Sv*). *WSC3* encodes for a cell wall integrity and stress response component (WSC) domain containing protein and is involved in hyphal fusion during plant colonization and fungus-fungus interactions (Wawra *et al*., 2019). Other commonly induced genes were LysM-domain chitin-binding proteins (e.g. the Pirin1_72967; PIIN_02170), chitin deacetylases (e.g. Pirin1_78560; PIIN_07753 in *Si* and Sebve1_327649 in *Sv*), fungal hydrophobins (Pirin1_77300; PIIN_06495 in *Si* and Sebve1_328749 in *Sv*), and secreted *Egh16-like* virulence factors (Pirin1_75341; PIIN_04536 in *Si* and Sebve1_ 330942 in *Sv*). Hydrophobins play a role in both plant colonization and mycoparasitic fungus-fungus interactions (Guzmán-Guzmán *et al*., 2017) and homologs of *Egh16-like* virulence factors are associated with cell wall remodelling and appressoria-mediated host penetration in plant, insect, and animal colonizing fungi (Xue *et al*., 2002; Grell *et al*., 2003; Cao *et al*., 2012; Herrera-Estrella *et al*., 2016; Huang *et al*., 2019; Shang *et al*., 2021), suggesting a dual function in plant colonization and mycoparasitism.

Additionally, we found several genes encoding for proteins with conserved DELD motifs (Lahrmann *et al*., 2013) to be upregulated *in planta* and during confrontation with *Bs*. Members of the DELD effector family, in particular *Dld1* (Pirin1_76679; PIIN_05872), have been shown to promote plant colonization by enhancing micronutrient availability to the fungus and interfering with oxidative stress and redox homeostasis (Nostadt *et al*., 2020). The function of DELD proteins during interfungal competition, however, remains to be functionally characterized.

### Sebacinales induce specific sets of effector candidates in response to host plants or microbes

To investigate whether mycoparasitism, microbial cooperation and colonization of plant hosts require the expression of secreted effector genes, we identified putatively secreted proteins in the Sebacinales with the Predector pipeline (1,183 in *Si* and 1,434 in *Sv*). A substantial share of effector genes (40 % in *Si* and 26 % in *Sv*) was significantly upregulated (>2 log2FC, adjusted p-value < 0.05) in response to at least one biotic interaction partner at one or more time points (Figure 2). The vast majority of these genes were specifically upregulated in response to plant hosts (50 % in *Si* and 66 % in *Sv*) or induced by both plants and microbes (42 % in *Si* and 26 % in *Sv*). A smaller proportion (8 % in both, *Si* and *Sv*) of the putative effector genes were specifically induced by microbes, encompassing small-secreted proteins (SSPs) and carbohydrate-active enzymes (CAZymes). The transcriptional activation of distinct sets of effector genes in the Sebacinales during plant or microbe interactions highlights the significance of plant-specific and dual responses to both plant and microbe.

**Figure 2:**
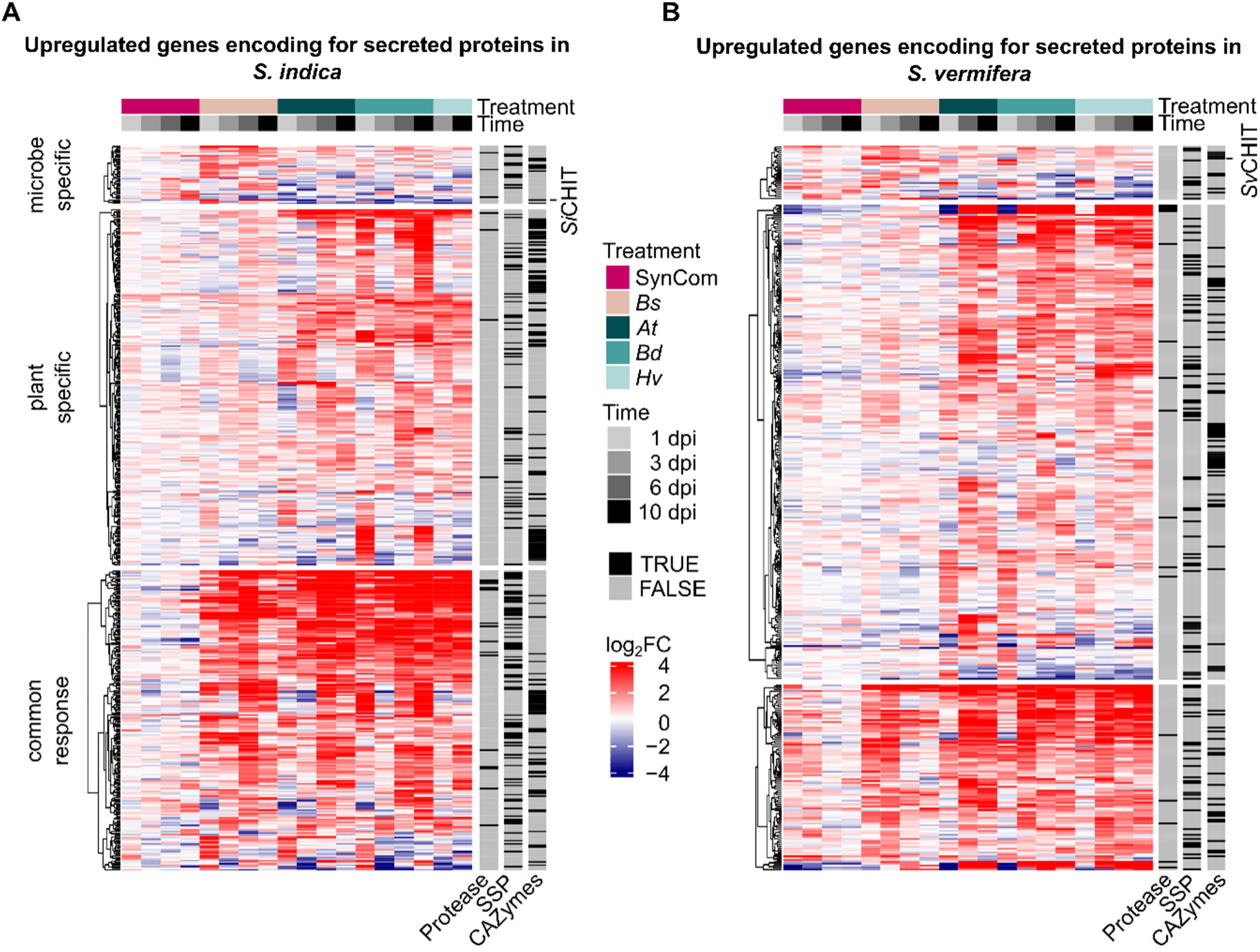
Interaction partner specific expression patterns of putative effector-coding genes in A) *S. indica* and B) *S. vermifera*. Genes encoding for secreted proteins were identified through the Predector pipeline (n = 1,183 for *Si* and 1,434 for *Sv*). Of these genes, 467 (*Si*) or 373 (*Sv*) were upregulated significantly (FDR-adjusted p-value < 0.05 and log_2_FC > 2) in response to at least one biotic interaction partner at one or more time points. These genes were annotated as “proteases” or “small secreted proteins” (< 300 amino acids) by the pipeline described by Pellegrin et al. (2015) or in case of “CAZymes” by the Predector pipeline. The clustering was performed separately for genes upregulated specifically in response to microbes (top), plants (centre) or both (bottom).

### The GH18-CBM5 chitinases are exclusive to the Basidiomycota within the fungal kingdom

One of the most prominently *Bs*-specifically upregulated gene in both *Si* and *Sv* was a chitinase from the GH18 family, with a CBM5 carbohydrate binding motif (Figure 3A). Fungal nutrient acquisition heavily relies on the secretion of CAZymes, particularly in breaking down soil organic matter and the cell walls of living plants and other fungi. Fungi across different divisions express an array of GH18 chitinases, each playing diverse roles in fungal development, nutrient uptake, and interactions with other organisms (Ihrmark *et al*., 2010; Chen *et al*., 2020). Here we investigated the distribution of GH18 chitinases across 135 distantly related fungal species spanning Ascomycota, Mucoromycota and Basidiomycota alongside different fungal lifestyles (Supplementary Figure 4). Within the fungal kingdom, chitinases with a GH18 domain featuring a CBM5 domain are solely present in the Basidiomycota (Figure 3B). The occurrence of GH18-CBM5 chitinases in Basidiomycota does not seem to be related to the fungal lifestyle, as they are found in saprotrophic as well as beneficial and phytopathogenic fungi and their copy number varies among species, suggesting functions in niche occupation or mycoparasitism rather than cell wall remodelling. Both *Si* and *Sv* carry only one copy of the GH18-CBM5 chitinase (Pirin1_74346; PIIN_03543 hereafter *Si*CHIT; Sebve1_16391, hereafter *Sv*CHIT) with an amino acid similarity of 78 %. This and the similar expression profiles of *Si*CHIT and *Sv*CHIT indicate a highly specialized conserved function of these chitinases in both Sebacinales (Figure 3).

**Figure 3:**
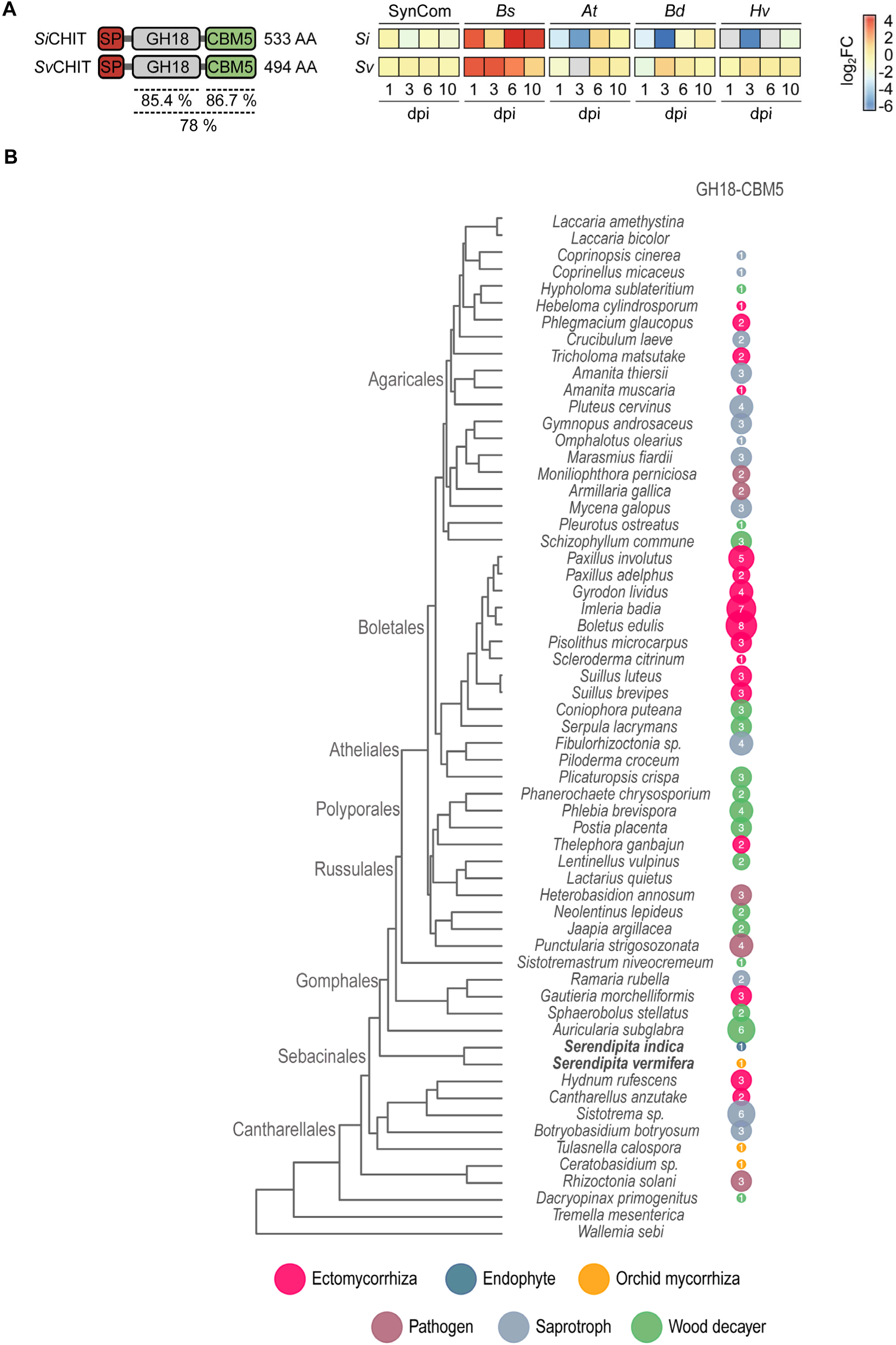
GH18-CBM5 chitinases are widespread among the Basidiomycota independent of their lifestyle. **A)** Domain architecture and expression pattern of *Si*CHIT and *Sv*CHIT during different biotic interactions at various time points. Percentages show the sequence similarity. **B)** Occurrence of GH18-CBM5 chitinases in different Basidomycota with varying lifestyles.

### *Si* reduces *Bs* infection and disease symptoms *in planta*

We previously reported that *Sv* mediates protection against *Bs* in barley and *At* and hypothesized that this protective function is linked to the secretion of antimicrobial effectors (Sarkar *et al*., 2019; Mahdi *et al*., 2022). Given the specific induction of *Sv*CHIT upon exposure of *Sv* to *Bs*, we speculated that the enzyme might contribute to antagonism between the beneficial root endophyte and the plant pathogen *Bs*. The conserved domain organization and expression pattern indicates that this might also be true for *Si*CHIT. To test this hypothesis, we first confirmed that *Si* displays a plant-protective phenotype against *Bs* in our host plants species (Li *et al*., 2023). To this end, we co-inoculated the roots of barley seedlings with *Si* and *Bs* spores and quantified fungal colonization by RT-qPCR six days post inoculation (dpi). We found that root colonization by *Bs*, but not *Si*, was drastically reduced in the co-inoculated roots compared to roots inoculated with only one fungus (Figure 4A). To assess disease symptoms, we measured root fresh weight and found that the reduced root colonization by *Bs* in the presence of *Si* correlated with a reduction of root growth inhibition (Figure 4B). Plant protection by *Si* was not linked to an increased expression of the barley defense marker gene *HvPR10* (Figure 4C), suggesting that the host protective capabilities of *Si* did not rely on an induction of plant immunity. To test whether the protective ability of *Si* from *Bs* was host-species independent, we assessed the fungal colonization (Figure 4D) and main root length (Figure 4E) of *At* seedlings upon co-inoculation with both fungi. In agreement with previous studies, we observed a growth-promoting effect of *Si* on the *At* seedlings in bipartite interactions in this host (Del Barrio-Duque *et al*., 2019; Scholz *et al*., 2023). In addition, the colonization of *At* by *Bs* and *Bs*-dependent reduction of root elongation were decreased in the presence of *Si.* We further monitored the progression of disease symptoms via Pulse-Amplitude-Modulation (PAM) fluorometry and demonstrated that co-inoculation with *Si* abolished the detrimental effects of *Bs* on photosynthetic activity in *At* leaves (Figure 4F). In summary, our findings demonstrate that *Si* has plant-protective abilities against the aggressive root rot pathogen *Bs* in barley and *At* but the molecular mechanisms that underlie this plant-protective ability are unknown.

**Figure 4:**
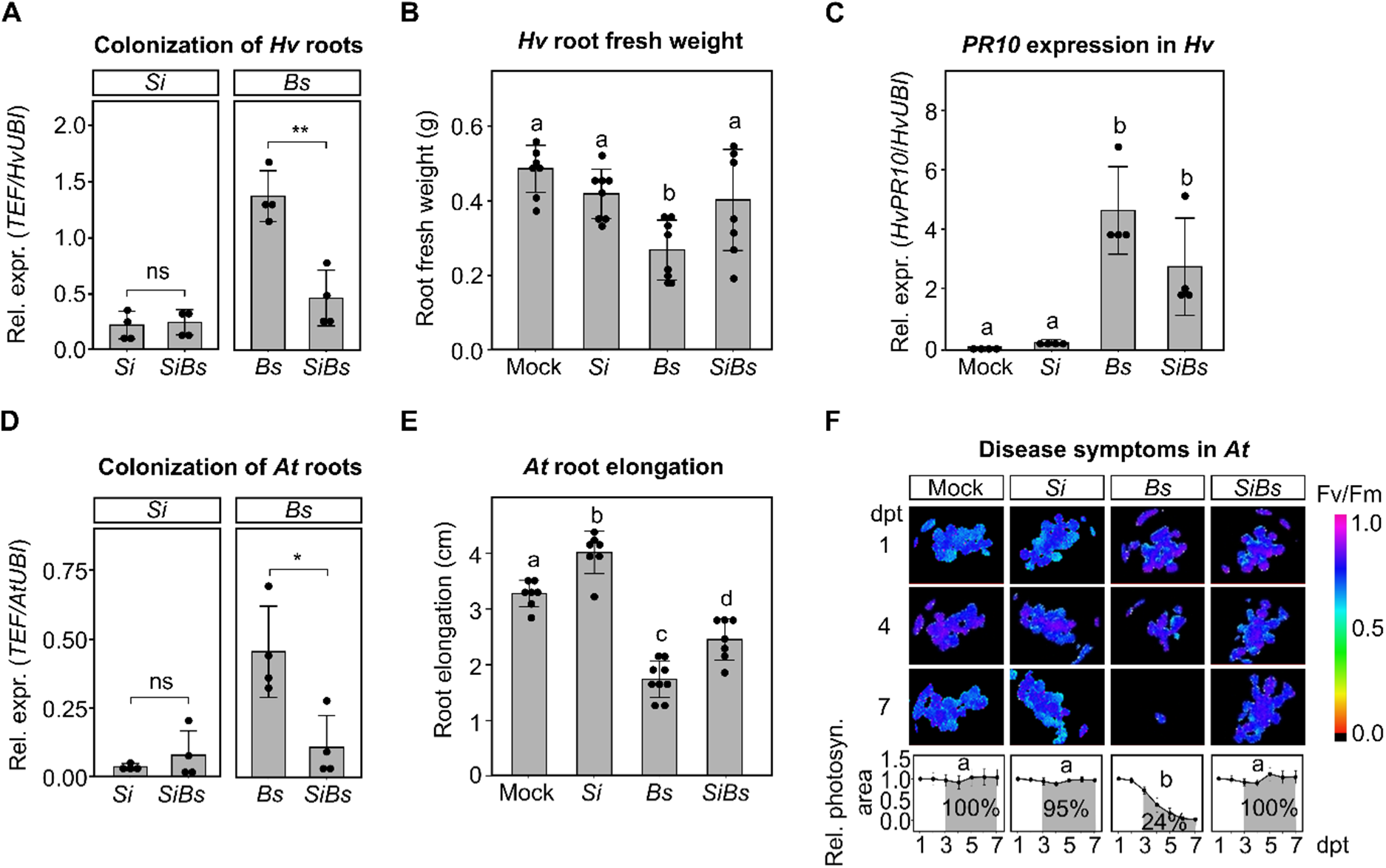
Plant-protective ability of *S. indica* in barley (A-C) and *Arabidopsis thaliana* (D-F). **A)** *Si* and *Bs* colonization at six dpi in barley roots inferred from the relative expression of the fungal housekeeping gene *TEF* compared to the barley ubiquitin (*HvUBI*) gene. For each replicate (n = 4), four plants were pooled. **B)** Barley root fresh weight after inoculation with *Si*, *Bs* or both fungi at six dpi. For each replicate (n > 7), four plants were pooled. **C)** *HvPR10* expression during mono- and co-inoculation of *Hv* with *Si* and *Bs* at six dpi. For each replicate (n = 4), four plants were pooled. **D)** *Si* and *Bs* colonization at six dpi in *At* inferred from the relative expression of the fungal housekeeping gene *TEF* compared to the Arabidopsis ubiquitin (*AtUBI*) gene. For each replicate (n = 4), ten plants were pooled. **E)** *At* root elongation at six dpi with *Si* and *Bs*. For each replicate, ten plants were pooled. **F)** Top: *At* photosynthetic activity (*F*_V_/*F*_M_) at 1, 4, and 7 days post transfer (dpt) corresponding to 7, 10, and 13 days post inoculation (dpi) with *Si*, *Bs*, or both fungi together. Bottom: Quantification of the photosynthetic area. Values were internally normalized to the first day of measurement. The percentages represent the remaining photosynthetic activity after the onset of disease symptoms normalized to the Mock control (area shown in grey). For each replicate (n = 4), ten plants were pooled. Statistical analysis: Student’s t-test (p-value < 0.01) for **A** and **D**; one-way ANOVA followed by Tukey‘s honest significant difference test (adjusted p-value < 0.05) for **B**, **C**, **E** and **F**. Different letters indicate significant differences. Individual biological replicates are represented as points; bars indicate averages ± standard deviation.

### The GH18-CBM5 chitinases have chitinolytic activity and inhibit *Bs* growth

Since we found that the *Si* and *Sv* GH18-CBM5 chitinases are specifically upregulated during *Bs* antagonism, we decided to characterize the function of these chitinases in fungus-fungus interactions. In a first step, we modelled the 3D structures of both enzymes using AlphaFold and docked a chitin octamer into the catalytic cleft that contains a conserved DxDxE motif required for catalysis (Figure 5A and Supplementary Figure 5A). In both cases, the substrate docked in close proximity to the DxDxE motif, with the *N*-acetyl group located near the second aspartate (D194 in *Si*CHIT and D171 in *Sv*CHIT) and the β-1,4 glycosidic bond located below the catalytically indispensable glutamate (E196 in *Si*CHIT and E173 *Sv*CHIT). This arrangement is in line with crystal structures of other GH18 chitinase - substrate complexes (van Aalten *et al*., 2001). To test the catalytic activity of *Si*CHIT and *Sv*CHIT, we expressed the recombinant proteins without signal peptide in *E. coli* and purified them from the supernatant of lysed bacterial cultures (Supplementary Figure 5B). Thin-layer chromatography revealed that both chitinases were active on crystalline crab shell chitin (Supplementary Figure 5C). To experimentally validate the importance of the DxDxE motif, we generated catalytically inactive mutants of both enzymes by exchanging the glutamate in the DxDxE motif with glutamine (*Si*CHIT^E196Q^ or *Sv*CHIT^E173Q^). This amino acid exchange has been shown to abolish chitinolytic activity without disrupting the chitin-binding ability in other GH18 chitinase-like effector proteins (Fiorin *et al*., 2018). A chitinase activity assay on chitin azure verified the loss of chitinolytic activity in *Si*CHIT^E196Q^ and *Sv*CHIT^E173Q^ (Figure 5B, Supplementary Figure 5D).

**Figure 5:**
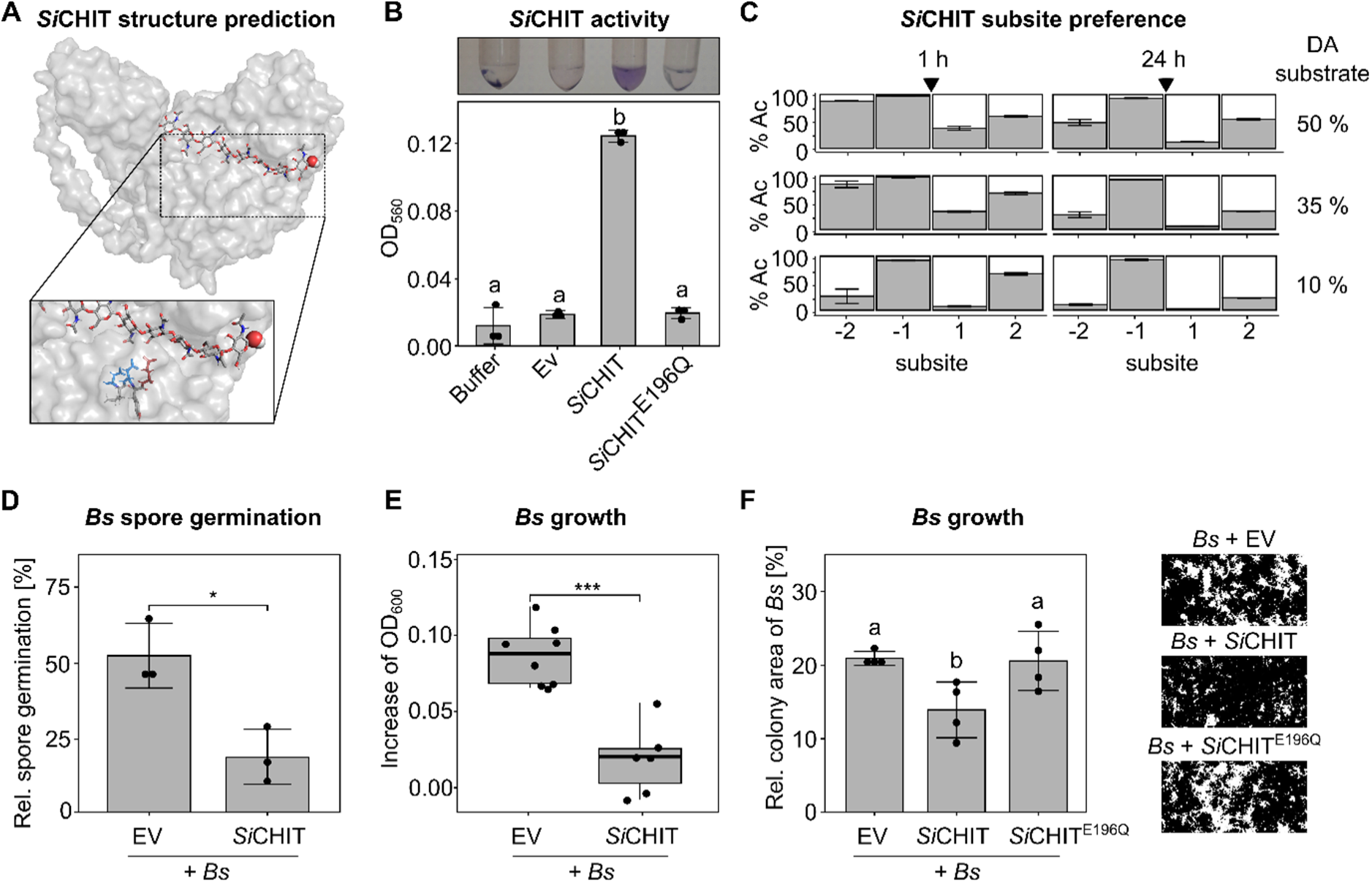
Recombinantly expressed *Si*CHIT is active and inhibits spore germination and growth of the plant pathogen *B. sorokiniana*. **A)** 3D structure of *Si*CHIT modelled using AlphaFold and visualized with PyMOL. The zoom-in shows the DIDYE motif, with aspartate (D) marked in blue and glutamate (E) marked in red. **B)** Chitinolytic activity of *Si*CHIT or the catalytically inactive *Si*CHIT^E196Q^. Chitin azure was incubated with 5 µM recombinant protein or the empty vector (Ev) control in 50 mM phosphate buffer (pH 6.0) for 24 h. Samples were spun down and the absorbance of the supernatant was measured at 560 nm (mean +/- SD, n = 3). **C)** Subsite specificity of *Si*CHIT as inferred by mass spectrometry. Chitosans of three degrees of acetylation (DA) were hydrolyzed for 1 h or 24 h and based on the sequenced products, the frequency of acetylated units at the -2 to +2 subsites of *Si*CHIT was determined. The black arrow indicates the glycosidic bond between the -1 and +1 subsite that is cleaved by the enzyme (mean +/- SD, n = 3). **D)** Relative *Bs* spore germination. Germinated and non-germinated *Bs* spores were counted 10 h after incubation with the recombinant chitinase or the Ev control. Student‘s t test was performed to determine statistical difference (*p < 0.05, mean +/- SD, n = 3) **E)** Increase of *Bs* OD_600_ 20 h after incubation with *Si*CHIT or the Ev control measured by OD_600_. Student‘s t test was performed to determine statistical difference (***p < 0.001, n = 6-8). **F)** Colony area of *Si*CHIT or *Si*CHIT^E196Q^-treated *Bs* six days after plating out on PNM medium. Left: quantification of *Bs* colony area. Different letters indicate significant differences (adjusted p-value < 0.05) according to one-way ANOVA followed by Tukey‘s honest significant difference (HSD) test (mean +/- SD, n = 4). Right: Exemplary pictures of *Bs* colonies treated with *Si*CHIT or *Si*CHIT^E196Q^.

GH18 chitinases exhibit a characteristic substrate specificity. They preferentially bind acetylated substrate units, but their -2, +1 and +2 subsites can also accept deacetylated substrate units, especially if the degree of acetylation of the substrate (DA) is low, such as with chitosan (Busswinkel *et al*., 2018). In contrast, the -1 subsite strictly requires acetylated units for catalysis. We examined the subsite specificity of *Si*CHIT by mass spectrometry of the oligomeric products generated during degradation of chitosan and found a substrate preference pattern consistent with the characteristic profile of GH18 chitinases (Figure 5C). Collectively, the thin-layer chromatography and the chitinase activity assay provide clear evidence that *Si*CHIT and *Sv*CHIT are typical GH18 chitinases which can degrade chitin and partially deacetylated chitosan *in vitro*.

To investigate the biological role of these chitinases during intermicrobial interactions, we determined the germination rate of *Bs* spores in the presence of both enzymes. Incubation with *Si*CHIT resulted in a significant 2.8-fold decrease in germination of *Bs* spores (Figure 5D). Similarly, although to a lesser extent, *Sv*CHIT reduced the germination of Bs spores by 1.7-fold (Supplementary Figure 5E). We also examined *Bs* growth after exposure to *Si*CHIT using spectroscopic analysis and found a significant reduction compared to the empty vector (Ev) control (Figure 5E). The growth of *Si* remained unaffected by *Si*CHIT, suggesting that the root endophyte is resistant to the effects of its own chitinase (Supplementary Figure 5F). Assessment of *Bs* fungal colony growth and morphology after treatment with *Si*CHIT, *Si*CHIT^E196Q^, or EV control on PNM medium showed a decrease in *Bs* growth specifically after *Si*CHIT pretreatment, confirming the necessity of *Si*CHIT’s chitinolytic activity for inhibition (Figure 5F). Our results strongly suggest that the fungal GH18-CBM5 chitinases have antimicrobial activity against the phytopathogenic fungus *Bs*. This finding prompted us to investigate whether the exogenous application of *Si*CHIT can alleviate the disease symptoms caused by *Bs in planta*.

### *Si*CHIT reduces disease symptoms of *Bs* in Arabidopsis and barley

We previously showed that Sebacinales predominantly safeguard the plant host through direct interactions among microbes occurring outside the root system (Sarkar *et al*., 2019; Mahdi *et al*., 2022). Therefore, to test the biocontrol ability of *Si*CHIT in barley, we inoculated host seedlings with *Bs* spores pre-treated with purified *Si*CHIT, the catalytically inactive *Si*CHIT^E196Q^ or the Ev control. Treatment of *Bs* spores with *Si*CHIT but not *Si*CHIT^E196Q^ reduced the colonization success of the pathogen (Figure 6A). Similarly, the reduction of root weight caused by *Bs* was significantly lower when the spores were pre-treated with *Si*CHIT, but not *Si*CHIT^E196Q^ (Figure 6B). In comparison to the Ev control, treatment with *Si*CHIT or *Si*CHIT^E196Q^ did not affect the expression of the barley defense-marker gene *HvPR10* triggered by *Bs* (Figure 6C). This implies that neither *Si*-mediated (Figure 4C) nor the *Si*CHIT-mediated protection of barley was linked to a significant induction of plant immunity.

**Figure 6:**
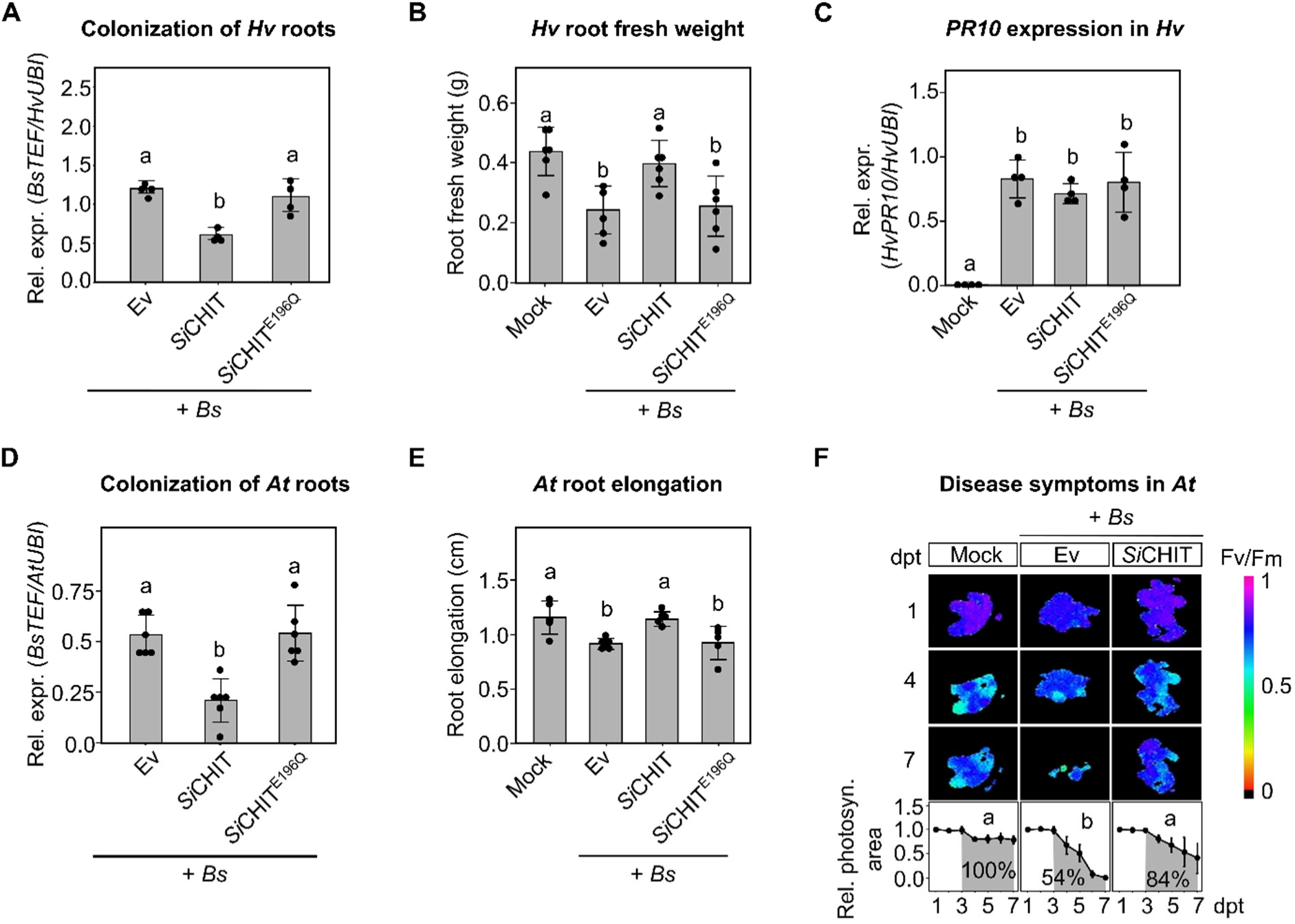
Plant-protective ability of *Si*CHIT in barley (A-C) and *Arabidopsis thaliana* (D-F). **A)** *Bs* root colonization at four dpi in barley roots inferred by from the relative expression of the fungal housekeeping gene *TEF* compared to the barley ubiquitin (*HvUBI)* gene. For each replicate (n = 6), four plants were pooled. **B)** Barley root fresh weight after inoculation with *Bs* spores, or *Bs* spores pre-treated with *Si*CHIT or *Si*CHIT^E196Q^ at four dpi. For each replicate (n = 6), four plants were pooled. **C)** *HvPR-10* expression in barley roots inoculated with *Bs* spores pre-treated with *Si*CHIT, *Si*CHIT^E196Q^ or the empty vector (Ev) control. For each replicate (n= 4), four plants were pooled. **D)** *Bs* colonization of *At* roots at four dpi with *Bs* spores, or *Bs* spores pre-treated with *Si*CHIT or *Si*CHIT^E196Q^ inferred from the relative expression of the fungal housekeeping gene *TEF* compared to the Arabidopsis ubiquitin (*AtUBI)* gene. For each biological replicate (n = 6), ten plants were pooled. **E)** *At* root length at four dpi with *Bs* spores, or *Bs* spores pre-treated with *Si*CHIT or *Si*CHIT^E196Q^. For each biological replicate (n = 5), ten plants were pooled. **F)** *At* photosynthetic activity (*F*_V_/*F*_M_) at one, four, and seven days post transfer (dpt) corresponding to seven, ten, and thirteen days post inoculation (dpi) with *Bs*, or *Bs* pre-treated with *Si*CHIT or *Si*CHIT^E196Q^. Bottom: Quantification of the photosynthetic area. Values were internally normalized to the first day of measurement. The percentages represent the remaining photosynthetic activity after the onset of disease symptoms normalized to the Mock control (area shown in grey). For each replicate (n = 4), ten plants were pooled. Statistical analysis: one-way ANOVA followed by Tukey‘s honest significant difference test (adjusted p-value < 0.05). Different letters indicate significant differences. Individual biological replicates are represented as points; bars indicate averages ± standard deviation.

Furthermore, we tested the plant-protective ability of *Si*CHIT in *At* by inoculating the seedlings with *Bs* spores pre-treated with *Si*CHIT, *Si*CHIT^E196Q^ or the Ev control. As previously observed in barley, root colonization (Figure 6D) and root growth inhibition (Figure 6E) by *Bs* were reduced when spores were treated with *Si*CHIT but not *Si*CHIT^E196Q^. To assess *Bs*-induced disease symptoms, we measured the photosynthetically active plant area over seven days via PAM fluorometry (Figure 6F). When *At* seedlings were treated with *Bs* and the Ev control, their cumulative photosynthetically active area from the onset of the first disease symptoms to the end of the experiment was reduced to 54 % of the mock control. Treatment of the *Bs* spores with *Si*CHIT resulted in a significantly lower reduction of the photosynthetically active area to 84 % of the mock control in the same time span. Similar to barley, this *Si*CHIT-mediated protection from *Bs* was not accompanied by an increased transcription of immune genes (Supplementary Figure 6).

These results demonstrate how the chitinolytic activity of *Si*CHIT reduces *Bs* viability, resulting in a significant decline in the pathogen’s ability to establish itself and cause harm to its host. This impact persisted throughout the entire experiment, confirming that host-independent intermicrobial interactions largely contribute to plant health in a complex tripartite system. Antimicrobial effectors like specialized chitinases emerge as pivotal players in multipartite interactions, contributing significantly to niche defense and beneficial effects by root endophytes.

## Discussion

In this study, we investigated the transcriptomic landscape of two closely related beneficial root endophytes in response to different host plants and root-associated microbes. We detected extensive transcriptional reprogramming in *Si* and *Sv* during host plant colonization and identified a set of genes that were commonly induced in the presence of all three host plants (*At*, *Bd* and *Hv*). These genes are likely to be general determinants of host colonization and are related to fungal nutrition, evasion or suppression of host immunity and, at later stages of the interaction, proteolysis and carbohydrate metabolic processes.

We have also identified host-species specific signatures underpinning colonization success. These different responses of *Si* and *Sv* to *At*, *Bd* and *Hv* are likely driven, at least in part, by divergences in host physiology and metabolism. For example, *Si* has been reported to respond to host-specific metabolic cues that directly affect the fungal colonization strategy. While *Si* establishes and maintains a biotrophic nutrition in living *At* epidermal cells, the endophyte switches rapidly to saprotrophy in response to nitrogen deficiency in *Hv* (Lahrmann *et al*., 2013). We also investigated the influence of the different cell wall composition of monocotyledons and dicotyledons on the response of Sebacinales to the three host plants. Our dataset confirmed the induction of genes encoding proteins specifically tailored for the degradation of monocotyledon cell walls, such as AA9, GH10 and GH11 (Lahrmann *et al*., 2013). The presence of such specialization underlines that cell wall degradation is important at certain stages of colonization. This observation is consistent with the broad host range of Sebacinales and the transition to a phase associated with cell death. Indeed, comparative genomics has shown that Sebacinales possess a higher number of CAZymes for plant cell wall degradation than merely biotrophic fungi, which are probably necessary to intracellularly colonize different host plants and thrive in soil ecosystems. Another factor underlying the host-specific transcription patterns in Sebacinales could be the different immune recognition of fungal-derived molecules by the hosts and the differences in the immune response. Taken together, these findings suggest that Sebacinales rely on highly complex and partially host-specific transcriptional profiles to effectively colonize a wide range of hosts. Such host-specific transcriptional responses have also been described in other polyspecialist fungi (Lahrmann *et al*., 2013; Lahrmann *et al*., 2015; Morán-Diez *et al*., 2015; Kusch *et al*., 2022).

In addition to the response of Sebacinales to host plants, we investigated the interaction of *Si* and *Sv* with a SynCom consisting of four taxonomically distinct bacterial strains from *At* roots. In co-culture with Sebacinales, these bacteria are tightly associated with the fungal glucan matrix and act synergistically with the root endophytes to mediate beneficial effects in both *At* and *Hv* (Mahdi *et al*., 2022). Despite the close physical interaction, root-associated bacteria elicited only minor transcriptomic changes in Sebacinales. This observation is consistent with the notion that most fungal responses to beneficial, neutral or antagonistic bacteria are attenuated within a few hours of initial contact (Mela *et al*., 2011; Deveau *et al*., 2015; Satterlee *et al*., 2022). Interestingly, however, a secreted GH32 invertase-like (Pirin1_79053; PIIN_08245 for *Si* and Sebve1_119217 for *Sv*) was upregulated in response to bacteria, other fungi and plants. This hints at the significance of breaking down sucrose into glucose and fructose during interactions across different kingdoms.

In contrast to the bacterial SynCom, the fungal competitor *Bs* elicited stronger transcriptomic changes in both *Si* and *Sv*. This response partially overlapped with the response to plant hosts, suggesting common underlying principles in the interaction of Sebacinales with plants and fungi. The secretion of effectors is a prime example of such a common principle. A substantial proportion of the differentially expressed putative effectors were induced in response to the presence of plant hosts and the phytopathogenic fungus *Bs*. We hypothesize that this set of core effectors may be involved in evading adverse responses from host plants or other microbes, possibly by shielding and remodeling their own fungal cell wall. Alternatively, these effectors could also serve as self-defense against hydrolytic enzymes and other toxic compounds secreted by plants or competing microbes (Snelders *et al*., 2018).

In addition to these core effectors, we have identified a number of effectors that are induced exclusively in response to host plants or root-associated microbes. We speculate that these groups may be enriched for effectors with phytosymbiotic or antimicrobial properties. Effector genes induced specifically in response to microbes accounted for only 8 % of all significantly induced effectors, supporting the hypothesis that soil-dwelling fungi constitutively express effectors with antimicrobial properties to deter competitors due to the ubiquitous presence of myriads of microbes in the soil (Snelders *et al*., 2018). In this scenario, expression of most antimicrobial effectors would not require specific cues. We furthermore found differences in the expression of glycosyltransferase CAZymes. These include a GT8, which is expressed consistently in all plant hosts but not in the presence of microbes, and a GT32, which is induced at a later stage of colonization and may therefore be involved in the establishment of long-lasting symbioses with the plant hosts. Despite the lower number of *Bs*-specifically expressed effectors, we identify and functionally characterize two Sebacinales GH18-CBM5 chitinases that were solely expressed in response to *Bs*. We demonstrated that these enzymes exhibit chitinolytic activity leading to antimicrobial activity against the plant pathogen *Bs*, but not against the Sebacinales themselves. The role of fungal GH18 chitinases during mycoparasitism and biocontrol has been extensively discussed for the Ascomycota fungi *Trichoderma* spp. (Carsolio *et al*., 1994; Woo *et al*., 1999; Druzhinina *et al*., 2011). These chitinases do not bear the CBM5 domain. GH18-CBM5 chitinases occur in bacterial taxa where the CBM5 domain is primarily relevant for substrate binding and degradation of crystalline chitin (Horn *et al*., 2006; Liu *et al*., 2023). Fusion of a bacterial-derived CBM5 with the *Trichoderma* GH18 chitinase results in enhanced substrate degradation and antagonistic activity toward other fungi (Limón *et al*., 2001; Limón *et al*., 2004), showing that the CBM5 can improve the performance of a biocontrol chitinase. Thus, the presence of a naturally occurring GH18-CBM5 chitinases in the Basidiomycota suggests that these fungi have evolved an effective strategy to combat other fungi. The mechanism by which *Si* and *Sv* protect themselves from the chitinolytic activity of the own GH18-CBM5 chitinase is not yet clear. However, it has been hypothesized that mycoparastic fungi limit the accessibility of their own cell walls to hydrolyses by expressing cell wall proteins that shield chitin while secreting an aggressive cocktail of enzymes designed to weaken the prey fungus (Gruber & Seidl-Seiboth, 2012). We have previously shown that various lectins are produced in *Si* and attach to the fungal cell wall and surrounding soluble glucan matrix, such as WSC- and LysM-domain containing proteins (Wawra *et al*., 2019), which are strongly induced upon confrontation with *Bs*.

A recent study highlighted that 75% of highly expressed CAZyme genes in ECM are linked to fungal cell wall degradation enzymes like AA5 (oxidases), GH5_9 (β-1,3-glucanases), GH18 (chitinases), GH20 (β-N-acetylglucosaminidases) and GH128 (β- 1,3-glucanases) families. Certain ECM species utilize GH18 and GH20 to break down chitin, suggesting that the degradation of fungal cell walls, especially chitin, by ECM plays an important role in the nitrogen cycle in forest soils (Lindahl & Taylor, 2004; Auer *et al*., 2023; Maillard *et al*., 2023). This mechanism might also apply to Sebacinales fungi. Since the genomes of Sebacinales are enriched for chitin and glucan binding lectins and GH18-CBM5 chitinases are widely distributed among Basidiomycota, this disease protection mechanism may be more widespread among endophytic fungi and ECM than previously thought (Govinda Rajulu *et al*., 2010). The secretion of a GH18-CBM5 chitinase may serve the nutritional needs of root-associated fungi through two strategies - by consuming the biomatter of the fungal competitor *Bs* and by safeguarding their ecological niche, the host plant, from the plant pathogen. This finding suggests that the effector-mediated manipulation of the microbiome by beneficial fungi extends beyond bacteria to fungal members of the plant microbiome.

## Material and methods

### Plant, fungal and bacterial materials

*Hordeum vulgare* (*Hv*, L. cv Golden Promise), *Brachypodium distachyon* (*Bd*, Bd21-3) and *Arabidopsis thaliana* (*At*, Col-0) were used as plant hosts. *Serendipita vermifera* (*Sv*; MAFF305830), *Serendipita indica* (*Si*; DSM11827) and *Bipolaris sorokiniana* (*Bs*; ND90Pr) were used as fungal models. The bacterial SynCom consists of four taxonomically diverse strains from the *At*Sphere collection (R11, R172, R189 and R935) which were described previously (Mahdi *et al*., 2022).

### Growth conditions and microbial inoculations

*Hv* and *At* seeds were sterilized and germinated as previously described (Mahdi *et al*., 2022). *Bd* seeds were sterilized in 3 % sodium hypochlorite and 0.1 % Triton-X for 30 min under constant shaking and then washed four times with sterile water every 15 min. Seeds were stratified for ten days in darkness at 4 °C on wet filter paper and subsequently transferred to sterile glass vials containing 1/10 PNM (Plant Nutrition Medium, pH 5.7) for germination on a day-night cycle of 16/8 h at 22/18 °C, 60 % humidity, and a light intensity of 108 µmol/m^2^s for eight days. *Sv* was propagated on MYP medium (Lahrmann *et al*., 2015), *Si* on CM medium (Hilbert *et al*., 2012) and *Bs* on modified CM medium (Sarkar *et al*., 2019), each containing 1.5 % agar, at 28 °C in darkness for 21 (*Si* and *Sv*) and 14 (*Bs*) days, respectively. Mycelial suspensions of *Sv* and spore suspensions of *Si* and *Bs* were prepared as previously described (Hilbert *et al*., 2012; Sarkar *et al*., 2019). Bacteria were grown in liquid TSB medium (Sigma Aldrich, St. Louis, USA, 15 g/l) at 28 °C in the dark at 220 rpm for 1-3 days depending on the growth rate. Bacterial suspensions were prepared as previously described (Mahdi *et al*., 2022). Plant roots were inoculated on 12 x 12 cm Petri dishes (*At*) or sterile glass jars (*Hv* and *Bd*) containing 1/10 PNM with *Sv* mycelium (0.12 g for *Hv* and *Bd* or 0.02 g for *At*), *Si* spores (500.000 chlamydospores per ml) or sterile water as control. Microbe-microbe confrontation experiments were performed on Petri dishes containing 1/10 PNM. Plates were inoculated with a) a pure suspension of *Sv* or *Si* mycelium (0.08 g), b) a mixed suspension of *Sv* or *Si* mycelium with *Bs* spores (10.000 spores) or c) a mixed suspension of *Si* or *Sv* with the bacterial SynCom (2 ml at OD_600_ = 0.01). All samples were kept on a day-night cycle of 16/8 hours at 22/18 °C, 60 % humidity, and 108 µmol/m^2^s light intensity for 1, 3, 6, and 10 days post inoculation (dpi). Samples for microbial confrontation were collected by scraping the fungal and bacterial material from the plate surface. Plant roots of all species were washed in MilliQ water to remove extraradical fungal hyphae. All samples were snap frozen in liquid nitrogen and used for RNA extraction. For plant protection assays in *At* or *Hv*, plants were co-inoculated with *Si* mycelium (0.02 or 0.12 g, respectively) and *Bs* spores (5.000 spores per plate or 15.000 spores per jar respectively). Plants were grown for six days before harvesting. Pulse amplitude modulation fluorometry (PAM) and ion leakage were used to assess disease symptoms in *At* (Mahdi *et al*., 2022). Fungal colonization was quantified in *At* and *Hv* by RT-qPCR using the primers listed in Supplementary Table S2.

### RNA extraction for RNA-seq analysis

RNA was extracted as previously described (Sarkar *et al*., 2019). RNA sequencing was performed at the U.S. Department of Energy Joint Genome Institute (JGI) under a project proposal (ID: 505829; Zuccaro 2020). Stranded RNA-seq libraries were generated and quantified by RT-qPCR. The sequencing was performed with Illumina technology in 151PE mode for each sample at the JGI. Raw reads were filtered and trimmed using the JGI QC pipeline. BBDuk was used to filter raw reads for artifact sequences by kmer matching (kmer=25), allowing one mismatch. Detected artifacts were trimmed at the 3’end. RNA spike-in reads, PhiX reads and reads containing NS were removed. Quality trimming was performed using phred trimming set at Q6. After trimming, the reads with a length below 25 bases or one third of the original read length were removed – whichever is longer. Filtered reads from each library were aligned to the *S. vermifera* MAFF 305830 v1.0 or *S. indica* DSM 11827 reference genomes downloaded from Mycocosm (https://mycocosm.jgi.doe.gov/mycocosm/home) using HISAT2 version 2.2.0. The raw gene counts were generated using featureCounts and the *Si* and *Sv* gff3 annotations. Only primary hits assigned to the reverse strand were included in the raw gene counts. To perform the PCA (principal component analysis), samples with low number of reads assigned to genes were not considered in the analysis. *Si* samples with more than 100,000 fragments assigned to genes were considered for the analysis. *Sv* samples with more than 46,000 fragments assigned to genes were considered with the exception of the AtSv1_1dpi, BdSv1_1dpi, BdSv2_3dpi samples. Subsequently, genes with less than a total of ten raw counts across all samples were filtered out. After the filtering, raw counts were normalised with the DESeq rlog transformations and PCA plot were drawn with the plotPCA function and customised with ggplot2.

### Differential gene expression analyses

The proportion of reads assigned to organisms per RNA-seq sample was examined. The consistency of normalized transcription for the biological replicates was confirmed by assessing the distribution of the number of genes and then the correlation of the biological replicates. Spearman’s rank correlation was calculated using the normalized number of genes of all biological replicates. Transcript counts of genes were normalized using the R package DESeq2 (Love *et al*., 2014) and then log_2_ transformed. Significant differentially expressed genes (DEGs) specific to conditions (> 2 log_2_FC; FDR-adjusted p < 0.05) were visualized using the R package UpSetR (Conway *et al*., 2017). Heatmaps were generated with the complexHeatmap package in R. K-means clustering was performed with the kmeans function in R setting the number of cluster to be generated to three. Functional annotations of the *S. indica* and *S. vermifera* genomes were downloaded from Mycocosm, Joint Genome Institute (https://mycocosm.jgi.doe.gov/mycocosm/home). Gene Ontology (GO) enrichment analysis was performed using the function enricher of ClusterProfiler setting the pvalueCutoff = 1.

### Fungal gene annotations

Secreted proteins and carbohydrate-active enzymes (CAZymes) were identified with Predector (Bendtsen *et al*., 2004; Sperschneider *et al*., 2016; Sperschneider *et al*., 2018a; Sperschneider *et al*., 2018b; Almagro Armenteros *et al*., 2019; Jones *et al*., 2021; Kristianingsih & MacLean, 2021; Sperschneider & Dodds, 2022; Teufel *et al*., 2022). Proteins with a manual secretion score > 0 and predicted as secreted were considered as secreted proteins. The protein domains of CAZymes were manually curated using the online tool dbCAN3 (https://bcb.unl.edu/dbCAN2/).

### Comparative analyses

Secreted CAZymes, proteases, and small secreted proteins (< 300 amino acids) of the fungi were identified (Pellegrin *et al*., 2015). We used the CAZyme family of GH18 conjugated with carbohydrate binding modules (CBMs) and the evolutionary order of the species from the comparative genomics of 135 fungi (Miyauchi *et al*., 2020). Output files generated above were combined and visualized with Proteomic Information Navigated Genomic Outlook (PRINGO).

### Chitinase expression and purification in *E. coli*

The coding sequences of *SiCHIT* and *SvCHIT* were amplified using the primers listed in Supplementary Table S1 and cloned into an expression vector (pQE-80L, Qiagen, Hilden, Germany) using Gibson assembly (New England Biolabs, Frankfurt, Germany). The resulting vector was transformed into chemically competent *E. coli* Mach1 cells. Protein expression was induced using 1 mM IPTG. The bacterial cultures were incubated at 16 °C and 120 rpm overnight. Bacteria were harvested via centrifugation at 5,000 g and 4 °C. For protein extraction, 6.5 ml lysis buffer (50 mM NaH_2_PO_4_, 300 mM NaCl and 10 mM imidazole) were added per g of bacterial pellet. Pellets were sonicated three times for 20 secs on ice and centrifuged at 13,500 rpm for 30 min. 5 ml of the supernatant were added to 1 ml of Nickel-NTA Agarose (Qiagen, Hilden, Germany) and incubated under continuous shaking at 4 °C for 1 h. Samples were spun down at 500 g for 20 secs, the flowthrough was removed, 4 ml wash buffer (50 mM NaH_2_PO_4_, 300 mM NaCl and 20 mM imidazole) were added and samples were incubated under continuous shaking at 4 °C for 10 min. Washing steps were repeated five times. For elution, 1 ml of elution buffer (50 mM NaH_2_PO_4_, 300 mM NaCl and 250 mM imidazole) was added and samples were incubated under continuous shaking at 4 °C for 10 min. Eluted fractions were dialyzed against 50 mM phosphate buffer (pH 6) over night and protein concentrations were measured using Bradford reagent. To test the purity of the protein samples, aliquots were boiled for 5 min in SDS-sample buffer, loaded on a 10 % SDS gel and visualized by Coomassie staining.

### Crab shell chitin hydrolysis and thin layer chromatography (TLC)

10 mg of crab shell chitin were weighed in 2 ml reaction tubes. Recombinant protein was added to a final concentration of 5 µM in 50 mM phosphate buffer. Samples were incubated at 28 °C over night. Next, insoluble chitin was spun down and 6.75 µl of the supernatant containing the hydrolysates was loaded on 60 F_254_ TLC silica gel plates (Sigma Aldrich, St. Louis, USA). The TLC was conducted as described previously (Li *et al*., 2002).

### Chitin azure assay

Chitin azure (Sigma Aldrich, St. Louis, USA) was adjusted to 4 mg/mL in 50 mM phosphate buffer (pH 6) and 100 µl were added to 2 ml reaction tubes. Recombinant protein was added to a final concentration of 5 µM in 200 µl. The samples were incubated at 28 °C and 120 rpm overnight. Next, samples were boiled at 95 °C for 5 min and centrifuged at 13,000 rpm for 10 min and supernatants were transferred to a 96 well plate. Absorption was measured at 560 nm.

### *Bs* spore germination assay

*Bs* spores were isolated as previously described (Sarkar *et al*., 2019) and diluted in TSB medium to a final concentration of 125,000 spores/ml. Recombinant protein was added to a final concentration of 5 µM, filled into 8 well chamber slides (VWR) and incubated for 6 h at 28 °C. The germination rate was quantified by non-invasive counting using an inverted microscope.

### *Bs* and *Si* growth assay

*Bs* and *Si* spores were isolated as previously described (Sarkar *et al*., 2019) and diluted in TSB medium to a final concentration of 125.000 spores/ ml. 200 µl spore stock were mixed with 200 µl protein extract to a final concentration of 5 µM, filled into 8 well chamber slides (VWR, Radnor, USA) and incubated for 6 h at 28 °C. The germination rate was quantified by non-invasive counting using an inverted microscope.

### *In planta* protection assays

To measure protection of *At* from *Bs*, *At* seeds were sterilized and germinated as described above. After transferring five-day-old seedlings to 1/10 PNM plates, they were inoculated with *Si* (500,000 spores), *Bs* (5,000 spores) or both fungi together, whereby *Si* was added two days earlier than *Bs*. To measure the protective role of *Si*CHIT, *Bs* spores were incubated in 50 mM phosphate buffer (pH 6.0) overnight with or without (Mock) 5 µM *Si*CHIT the day before plant inoculation. Five days after inoculation with *Bs*, seedlings were transferred into 24-well plates with water and PAM was measured over seven days (Dunken *et al*., 2022). For *Hv* inoculation, *Bs* spores were treated with recombinant enzyme as described above prior to root inoculation. After six days, *Hv* plants were harvested and roots were weighed after washing. Colonization of *Bs* was assessed by RT-qPCR following RNA extraction and cDNA synthesis as described previously (Sarkar *et al*., 2019).

### Docking

The 3D structure files of the chitin oligomer ligands (tetramer, hexamer, octamer) were created using the GLYCAM Web carbohydrate builder (Woods Group. (2005-2023) GLYCAM Web. Complex Carbohydrate Research Center, University of Georgia, Athens, GA. (http://glycam.org)), the models of *Si*CHIT and *Sv*CHIT without signal peptides were generated with AlphaFold (Jumper *et al*., 2021; Varadi *et al*., 2022). Amino acids within the models with a confidence measure pLDDT < 50 were identified and removed via PyMOL (Schrödinger, L. 2020 The PyMOL Molecular Graphics System, Version 2.5.), and the remaining two protein parts were split into two files, one for the catalytic core enzyme, one for the CBM5 domain. The scripts prepare_ligand4.py and prepare_ receptor4.py from the AutoDockTools (Morris *et al*., 2009) were used to prepare the ligand and receptor files before docking was performed with AutoDock VinaCarb v1.0 (Nivedha *et al*., 2016). Docking results were visualized and evaluated using PyMOL Version 2.5. (Schrödinger, L., & DeLano, W. 2020. PyMOL. Retrieved from http://www.pymol.org/pymol).

### Chitinase subsite specificity

The subsite specificity of *Si*CHIT was performed as previously described (Cord-Landwehr *et al*., 2017). *Si*CHIT was used at a final concentration of 0.2 µM on 1 mg/ml of chitosan with 10 %, 35 % or 50 % of acetylation in 50 mM of sodium acetate buffer. Chitosan oligomers produced by *Si*CHIT were quantified based on their degree of polymerization (DP) and degree of acetylation (DA) using HILIC-ESI-MS detection. The analysis was conducted using a Waters ACQUITY Premier UPLC System coupled to a Waters Synapt XS HDMS 4k mass spectrometer. The chitosan oligomers were N-acetylated using [2H^6^] acetic anhydride and then mixed with the internal double isotopic labeled R*1-6 standard. The oligomer mixtures were separated using an Acquity UPLC BEH Amide column (1.7 μm, 2.1 mm × 50 mm; Waters Corporation, Milford, MA, USA) and a VanGuard precolumn (1.7 μm, 2.1 mm × 5 mm; Waters Corporation, Milford, MA, USA). The flow rate was set to 0.4 ml/min, the column oven temperature was set to 40 °C, and 2 µl of the sample, including 50 ng of the internal standard (double isotopic labelled chitin oligomers DP 1-6), were injected into the system. The sample was separated using an LC run lasting 10.5 minutes, with the following gradient elution profile. 100 % A (80:20 ACN/H_2_O with 10 mM NH_4_HCO_2_ and 0.1 % (v/v) HCOOH) for 0-1 minutes, followed by a linear gradient to 20% (v/v) B (20:80 ACN/H_2_O with 10 mM NH_4_HCO_2_ and 0.1% (v/v) HCOOH) from 1.0 to 7.5 minutes, and then to 75 % (v/v) B from min 7.5 to 8.5 minutes. From minutes 8.5 to 9.0 isocratic 20 % B, followed by a linear gradient from 20 % (v/v) A to 100 % (v/v) A from minute 9.0 to minute 9.2. This was followed by an isocratic elution of 100 % A from minute 9.2 to minute 10.5. The MS1 measurements were conducted in positive resolution mode with normal dynamic range sensitivity, with a mass range of m/z 50 to 2000 and a scan time of 0.25 seconds. To improve mass accuracy, lock spray calibration was performed using leucine encephalin, injected in 10-second intervals for 1 second as a look mass. The capillary voltage of the source was set to 3 kV, the sampling cone to 35, the source offset to 20, while the source and desolvation temperatures were set to 80 °C and 250 °C, respectively. The cone gas flow was set to 0 l/h and the desolvation gas flow was set to 500 l/h. The nebulizer pressure was set to 5.8 bar. Quantitative pattern determination was performed following the method described in Cord-Landwehr et al. 2017 with the following modification. The LC-MS system used for the MS1 analysis was also used for this analysis. The TOF MS/MS mode was employed to determine the pattern of isotopically labelled and 18O-labelled oligomers. Each oligomer containing at least one natural and one isotopically labelled acetyl group was individually isolated using the quadrupole of the MS system with a LM Resolution of 15 to enable the isolation and fragmentation of mono-isotopic peaks. The collision energy in the trap was set to 18 V for dimers, 20 V for trimers, 22 V for tetramers, 24 V for pentamers and 26 V for hexameric products.

## Acknowledgements

We acknowledge the work (proposal: **10.46936/10.25585/60001292**) conducted by the U.S. Department of Energy Joint Genome Institute (https://ror.org/04xm1d337), a DOE Office of Science User Facility, is supported by the Office of Science of the U.S. Department of Energy operated under Contract No. DE-AC02-05CH11231. We thank the Cluster of Excellence of Plant Science (CEPLAS) and the SPP2125 “Decrypt” for financial support. We want to thank Prof. Dr. Francis Martin, Dr. László Nagy and Prof. Dr. Joseph Spatafora for providing the genome sequences generated in the framework of the 1000 Fungal Genomes (1KFG) project and Prof. Dr. Francis Martin for the pipelines for comparative analysis of the GH18-CBM5 chitinases.

