## Supplemental figures for "Time-resolved transcriptomics reveal a mechanism of host niche defense: beneficial root endophytes deploy a host-protective antimicrobial GH18-CBM5 chitinase"

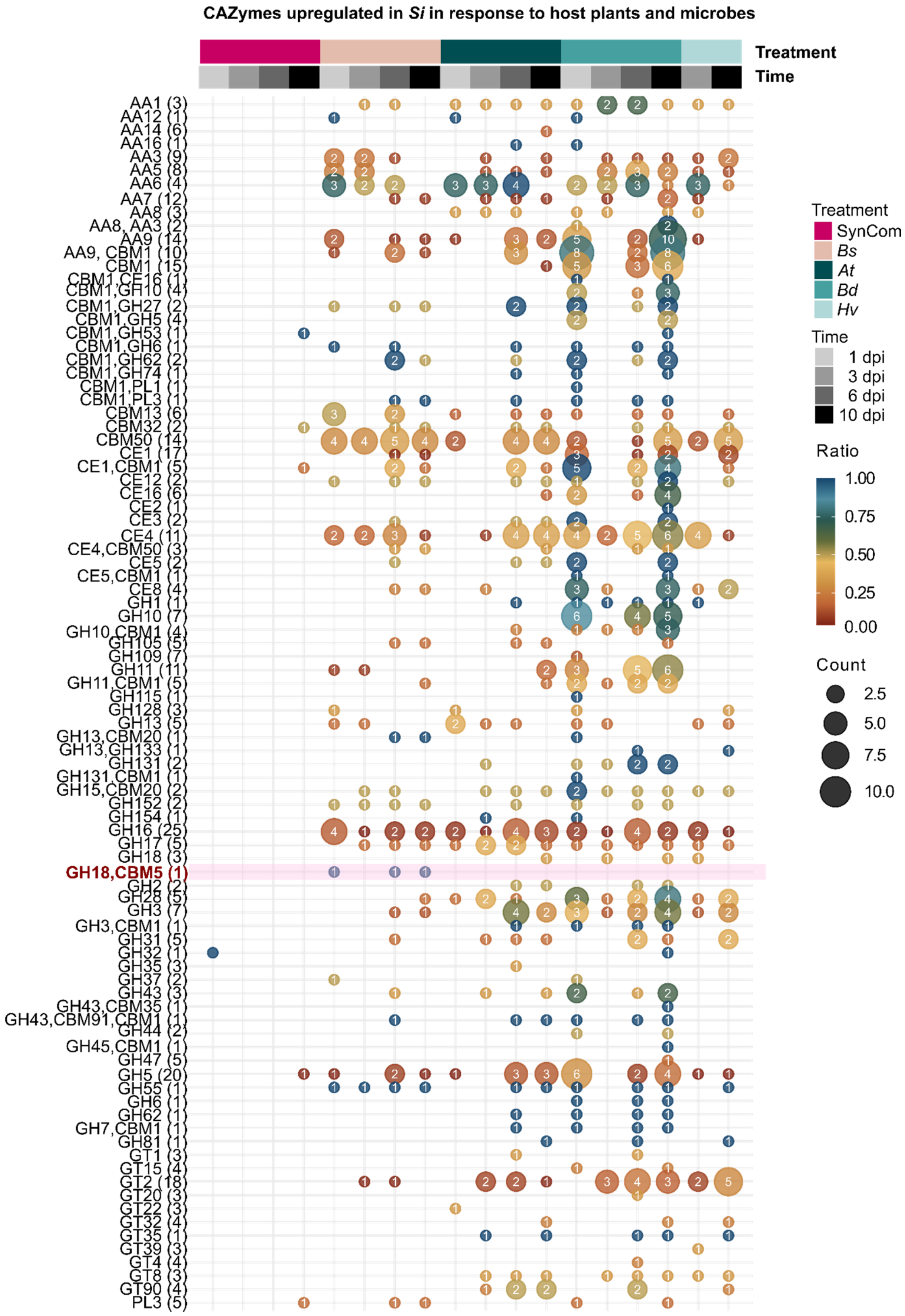


**Supplementary Figure S1: Upregulated *Si* CAZymes in response to host plants and microbes.** The bubble plot shows the protein domains of the CAZymes annotated in the genome of *Si* and up-regulated in at least one comparison. Numbers in each bubble indicate the number of up-regulated CAZymes with the indicated domain for each comparison. Colors indicate the ratio between the number of up-regulated CAZymes and total number (in brackets) of CAZymes with the considered domain in the genome. The CAZymes were annotated with the Predector pipeline and the protein domains were manually curated using the dbcan website. The GH18-CBM5 chitinase is highlighted in red.

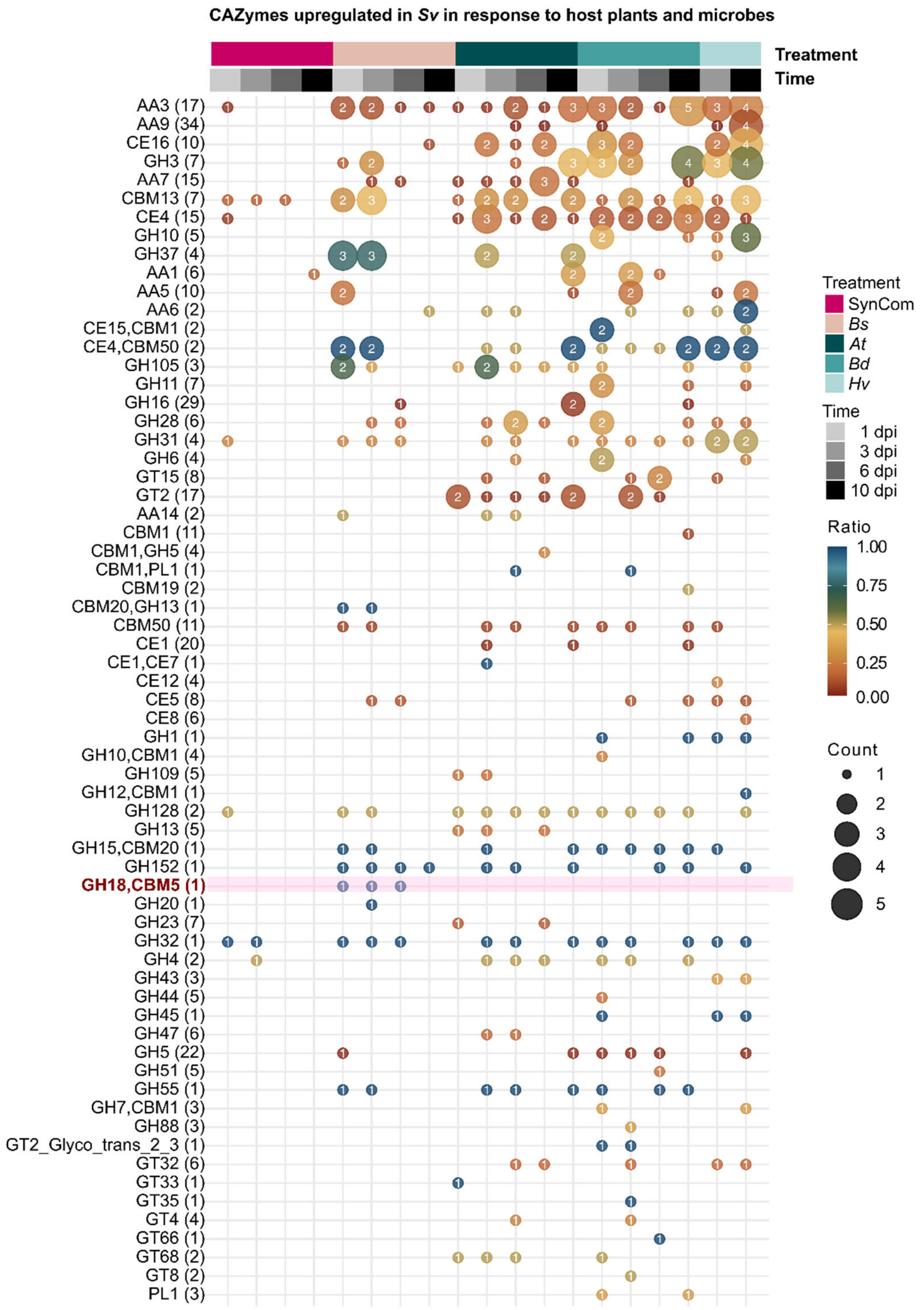


**Supplementary Figure S2: Upregulated *Sv* CAZymes in response to host plants and microbes.** The bubble plots shows the protein domains of the CAZymes annotated in the genome of *Sv* and up-regulated in at least one comparison. Numbers in each bubble indicate the number of up-regulated CAZymes with the indicated domain for each comparison. Colors indicate the ratio between the number of up-regulated CAZymes and total number (in brackets) of CAZymes with the considered domain in the genome. The CAZymes were annotated with the Predector pipeline and the protein domains were manually curated using the dbcan website. The GH18-CBM5 chitinase is highlighted in red.

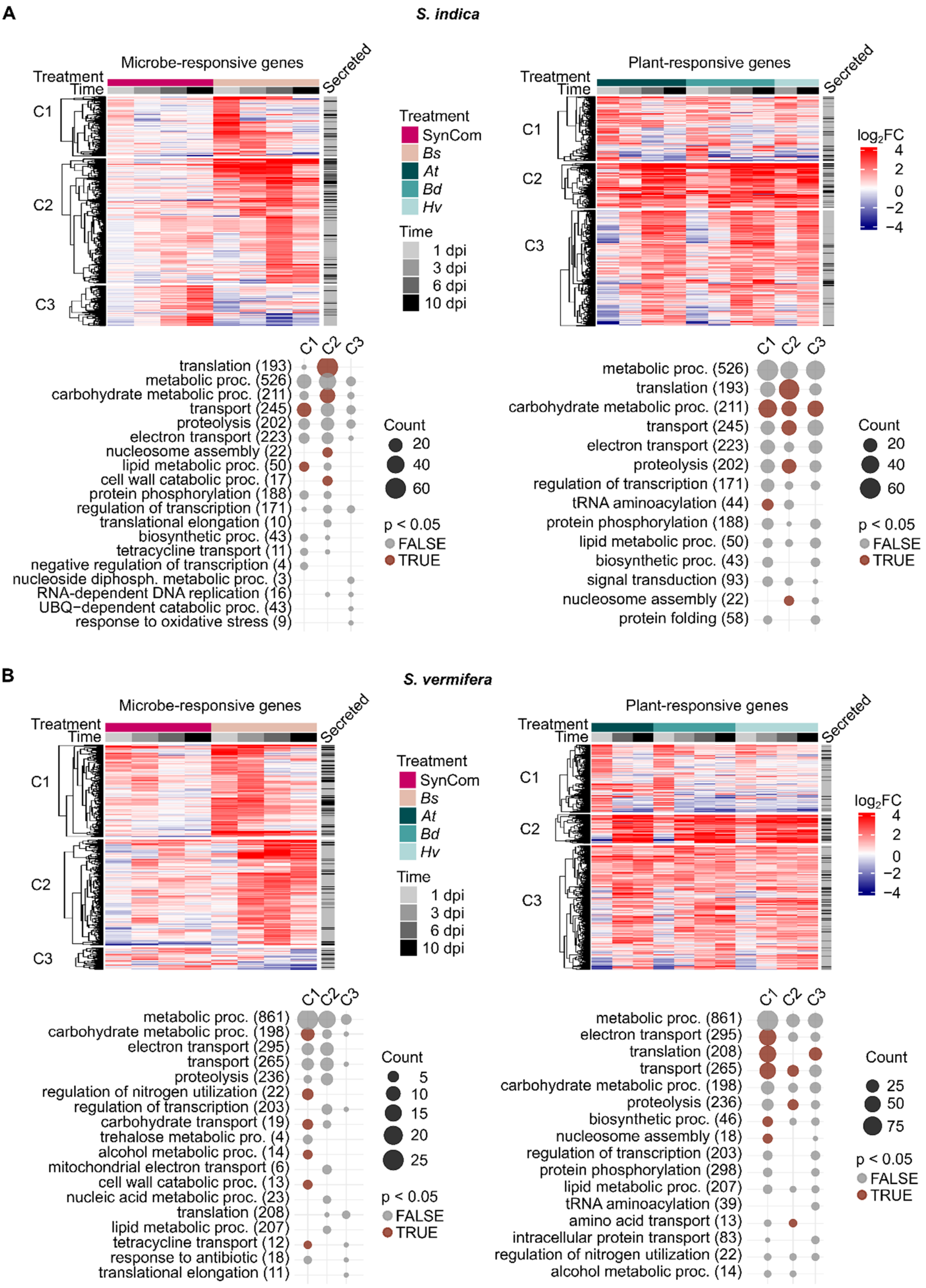


**Supplementary Figure S3: K-means clustering and Gene Ontology (GO) term annotation of microbe-responsive (left) and plant-responsive (right) upregulated genes in *S. indica* (A) and *S. vermifera* (B).** The GO term plots show the top ten most abundant biological processes for each cluster formed by the k-means clustering. The size of the dots indicates the number of genes associated with a specific GO term, whereas the colour indicates if the enrichment is significant (adjusted p-value < 0.05).

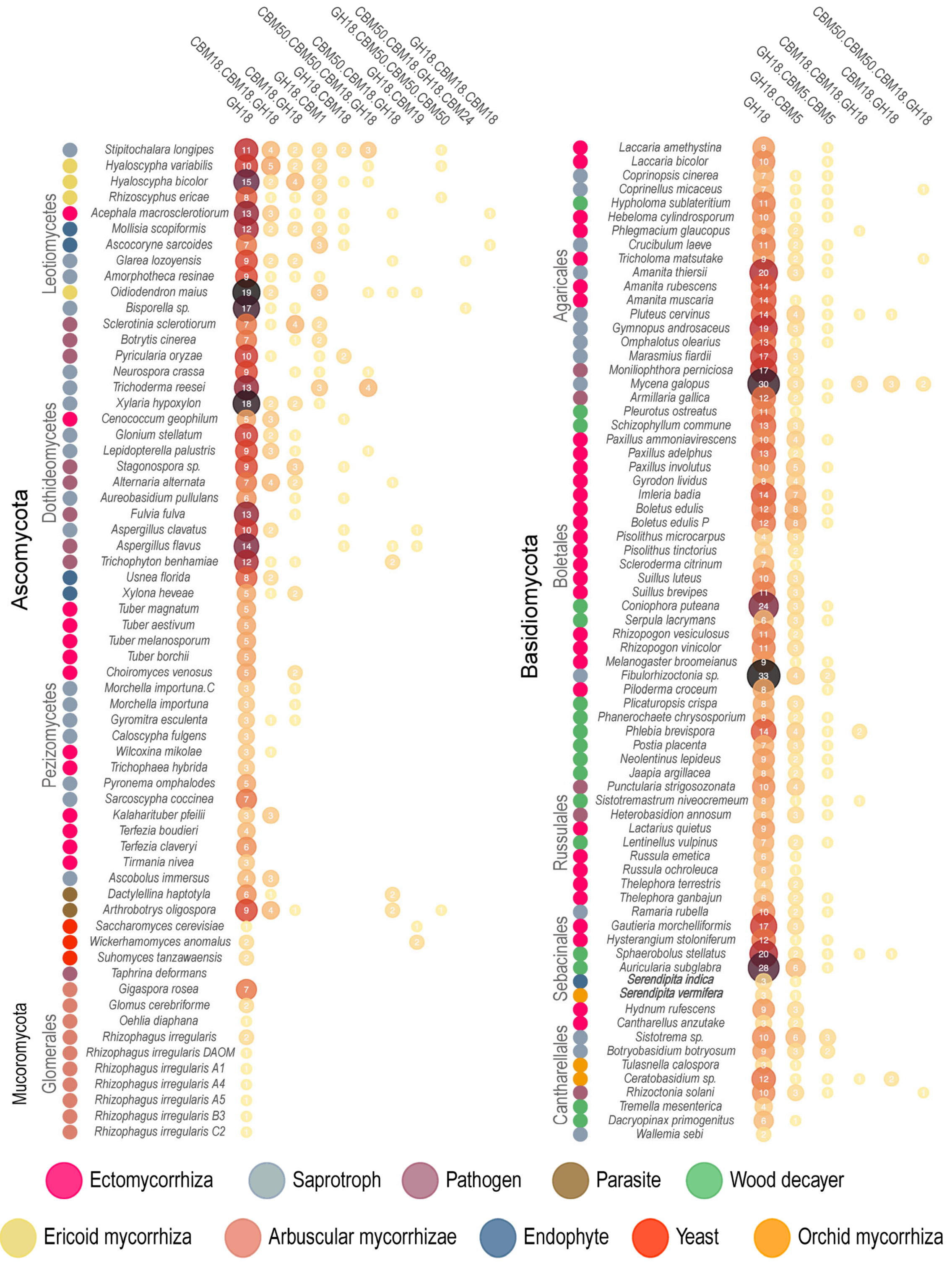


**Supplementary Figure S4: Occurrence of GH18 and different CBM domains in 135 fungi in three phyla.** Bubbles contain the number of genes encoding a GH18-conjugated CBM or a GH18 enzyme without a CBM. Taxa are color-coded according to their mode of life (see bottom panel). The evolutionary order of fungi was taken from the JGI mycocosm. Ascomycota and Mucoromycota (left) and Basidiomycota (right). No GH18-CBM5 is present in Ascomycota and Mucoromycota. The GH18-CBM5 column is highlighted in red. All genomes (Supplementary Table S3) have been published (Miyauchi *et al.*, 2020) or principal investigators have granted the usage.

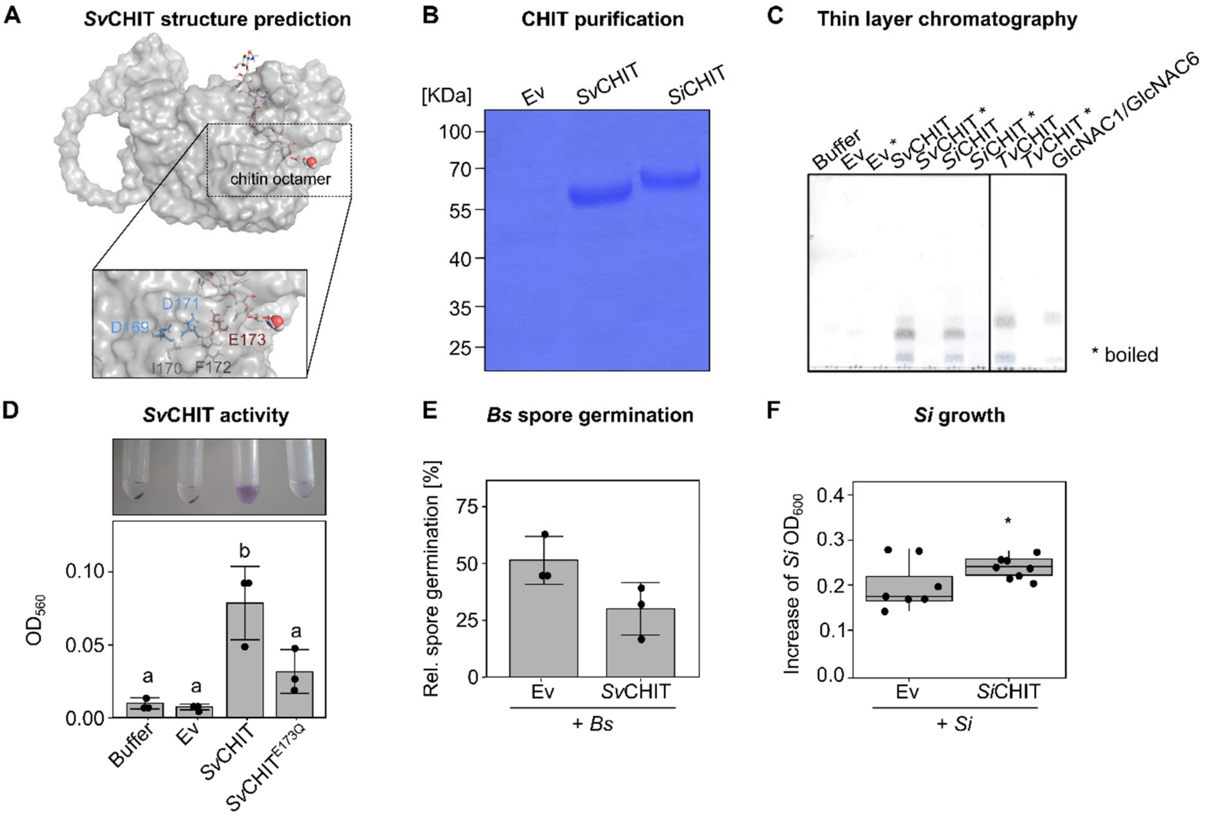


**Supplementary Figure S5: Purification of *Si*CHIT and *Sv*CHIT, *Sv*CHIT activity and effect of *Si*CHIT on *S. indica* growth. A)** 3D structure of *Sv*CHIT modelled using AlphaFold and visualized by PyMOL. The zoom-in shows the DIDYE motif, with aspartate (D) marked in blue and glutamate (E) marked in red. **B)** Purification of chitinases. *Si*CHIT and *Sv*CHIT were expressed in *E. coli*, purified via an N-terminal His-tag and visualized on a 10 % SDS-gel stained with Coomassie brilliant blue. **C)** Thin layer chromatography of crab shell chitin hydrolysates. Crab shell chitin was incubated with recombinant enzymes at 5 µM for 24 h at 37 °C in 50 mM phosphate buffer (pH 6). A chitinase mix from *Trichoderma viride* was used as positive control. **D)** Chitinolytic activity of *Sv*CHIT and the catalytically inactive *Sv*CHIT$^{E173Q}$ on chitin azure. Chitin azure was incubated with 5 µM of respective recombinant protein or the Ev control in 50 mM phosphate buffer (pH 6) for 24 h. Samples were spun down and the absorbance of the supernatant was measured at 560 nm. Individual biological replicates (n = 3) are represented as points; bars indicate averages ± standard deviation. **E)** Relative *Bs* spore germination. Germinated and non-germinated *Bs* spores were counted 10 h after incubation with the recombinant chitinase or the empty vector (Ev) control. **F)** Effect of purified *Si*CHIT on *S. indica* growth. *Si* spores were treated with 5 µM *Si*CHIT or the empty vector (Ev) control in ½ TSB medium and growth was assessed by $OD_{600}$ measurement. Data show OD values at 20 h compared to 0 h. Statistical difference was assessed by Student's t-Test (n = 7-8).

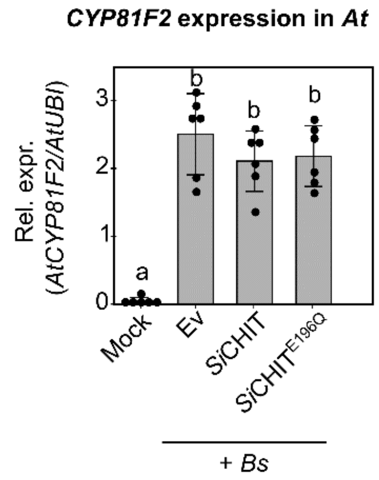


**Supplementary Figure S6: Defence gene expression after treatment of *At* seedlings with *Bs* pre-treated with *Si*CHIT.** *AtCYP18F2* expression in *At* roots inoculated with *Bs* spores pre-treated with *Si*CHIT, *Si*CHIT$^{E196Q}$ or the Ev control. Different letters indicate significant differences according to a one-way ANOVA followed by Tukey's honest significant difference (HSD) test (adjusted p-value < 0.05, mean +/- SD, n = 6).
